# How endosomal PIKfyve inhibition prevents viral membrane fusion and entry

**DOI:** 10.1101/2025.11.25.690586

**Authors:** Nicholas Chow, Gustavo Scanavachi, Anand Saminathan, Tom Kirchhausen

## Abstract

Enveloped viruses enter cells by membrane fusion. The viral membrane fuses with a host membrane, either at the cell surface or within endocytic compartments. For endocytic entry, fusion is typically triggered by low pH and often requires proteolytic priming by compartment-specific host proteases, which together define the site and mechanism of fusion and shape viral tropism.

Inhibition of the lipid kinase PIKfyve, which generates PI(5)P and PI(3,5)P₂ in late endosomes and lysosomes, swells those compartments and blocks infection by a subset of enveloped viruses, including Ebola virus, Marburg virus, coronaviruses (SARS-CoV-2), and VSV chimeras bearing Ebola, SARS-CoV-2, or Lassa glycoproteins, while showing minor effects on H1N1 influenza and no effect on VSV or VSV–rabies chimeras. In the work reported here, we have determined the basis for selectivity.

We show that swelling of late endosomes/lysosomes, independent of changes in lipid composition or altered virion trafficking, is sufficient to block virus–endosome fusion and genome release, even when endosomal acidity is preserved. Acute PIKfyve inhibition with apilimod or brief hypotonic treatment produced endosomal swelling and impaired infection by interrupting a late endosomal entry step. Live-cell 3D lattice light-sheet fluorescence microscopy imaging tracked fluorescent virions accumulating and arresting in late endosomes prior to fusion, and single-cell, single-round assays confirmed loss of infectivity.

These data support a simple biophysical mechanism: endo-lysosomal swelling, likely increasing endosomal membrane tension, creates an energy barrier to fusion and genome release. Inducing such swelling may offer a general strategy to inhibit viruses that depend on late endosomal entry.

**SIGNIFICANCE STATEMENT:** Why does inhibiting the endosomal lipid kinase PIKfyve, which generates PI(5)P and PI(3,5)P_2_, block entry of some enveloped viruses but spare VSV-G? Using single-round infectivity and live-cell 3D imaging, we show that acute PIKfyve inhibition traps incoming VSV chimeras bearing SARS-CoV-2 spike or Ebola GP in swollen late endosomes/lysosomes before fusion and genome release. Hypotonic swelling phenocopies the block, and removing glutamine prevents apilimod-induced swelling and largely restores infection, arguing that swelling—not altered phosphoinositides—restricts entry. We propose a physical mechanism: increased endosomal membrane tension raises the energetic barrier for fusion-pore formation and genome release. Modulating endolysosomal volume may therefore inhibit viruses that rely on late endosomal entry.

## INTRODUCTION

Enveloped viruses initiate infection by fusing their membrane with a host-cell membrane, delivering their genetic material into the cytosol (see review (1)). This fusion can occur at the plasma membrane or within endocytic compartments. For many viruses, including filoviruses and coronaviruses that rely on endocytosis, fusion is typically triggered by low endosomal pH and often requires proteolytic priming of viral glycoproteins by endosomal resident host proteases; for some, an additional host factor in the endolysosomal compartment is required (see reviews (2–5)). These spatial and pH dependencies define the site and mechanism of fusion during the early steps of infection, and, ultimately, determine viral tropism and infectivity.

PIKfyve, a lipid kinase on late endosomal and lysosomal membranes, generates phosphatidylinositol 3,5-bisphosphate [PI(3,5)P₂] and phosphatidylinositol 5-phosphate [PI(5)P₂] (6). Its depletion or pharmacological inhibition produces enlarged late endosomes and lysosomes and decreases PI(5)P and PI(3,5)P_2_ while increasing PI3P levels by preventing its conversion into PI(3,5)P_2_ (7–10); the organelle specificity of these alterations has been confirmed by the localization of optical phosphoinositide biosensors by fluorescence microscopy (11). PIKfyve inhibition blocks infection by a subset of enveloped viruses at early stages of entry. For example, treatment with the PIKfyve inhibitor apilimod prevents GP-mediated entry of filoviruses and SARS-CoV-2 and for VSV- or lentivirus-based pseudotypes, without affecting virion uptake or bulk endosomal acidification (12, 13). Vacuolin-1, which also targets PIKfyve, similarly suppresses single-cycle infection by VSV chimeras bearing Ebola or SARS-CoV-2 glycoproteins by preventing fusion and genome delivery from endolysosomal compartments (13). Other PIKfyve-directed inhibitors, including YM201636, SGC-PIKFYVE-1l and PIKfyve/PIP5K1C, inhibit early infection by parental African swine fever virus, enteroviruses, and β-coronaviruses, as well as by SARS-CoV-2 spike–pseudotyped virions (14–17). Together, these observations support the view that PIKfyve-activity at late endosomes and lysosomes is a common requirement for successful viral entry. In contrast, PIKfyve inhibition has only modest effects on H1N1 influenza virus infection (18) and little or no effect on viruses whose entry occurs in early endosomes, such as VSV and VSV particles bearing rabies virus G (VSV–RABV) (13).

A recent study identified LYVAC (PDZD8) as the ER-anchored lipid transfer protein that mediates ER-to-lysosome lipid transfer at ER–endolysosome contact sites, enabling osmotic membrane expansion (19). In the absence of LYVAC, apilimod fails to induce lysosomal swelling, supporting a model in which vacuolation primarily and the consequent increase in membrane tension depends on lipid transfer, rather than on loss of PI3P or PI(3,5)P₂. We therefore tested the hypothesis that the antiviral effect of PIKfyve inhibition by apilimod arises primarily from a mechanical constraint on virion fusion to the limiting membranes of the endolysosomal system, rather than from a block in endosomal trafficking or a change in endosomal phosphoinositide composition. We combined quantitative infectivity assays with high-precision, live-cell three-dimensional fluorescence microscopy to track single virions, allowing direct comparison of the effects of apilimod on VSV, VSV–ZEBOV, and VSV–SARS-CoV-2 infection.

Because these viruses share an identical VSV backbone, differences in apilimod sensitivity must arise from how their entry steps respond to perturbation of PIKfyve activity rather than from downstream replication events. Entry of VSV is mediated by its G protein and requires no further proteolysis for activation. Following clathrin-mediated endocytosis, acidification in early and early-late endosomes triggers a conformational change in G that drives membrane fusion (20). In contrast, VSV–ZEBOV requires two sequential processing steps of its GP: cleavage during biosynthesis, followed by cathepsin-mediated trimming in late endosomes or lysosomes to expose the NPC1-binding site. VSV–SARS-CoV-2 likewise depends on sequential proteolysis of its spike protein, furin cleavage at S1/S2 during maturation, followed by TMPRSS2- or cathepsin-mediated cleavage at S2′ and on endosomal acidification (see review (21)).

We show here that endosomal swelling, induced either by apilimod or by hypoosmotic challenge, correlates with inhibition of viral fusion, but only for VSV–ZEBOV or VSV–SARS-CoV-2 virions that reach and attempt fusion within swollen compartments. VSV, which enters from early endosomes that do not swell, escapes this inhibition, as does a subpopulation of VSV–SARS-CoV-2 virions that fuse in early endosomes. We suggest that under these swelling conditions, endolysosomes accumulate solute and water and increase their volume without a fully compensatory increase in membrane area. The resulting rise in membrane tension will oppose the local membrane remodeling required for fusion (1). Increased membrane tension in swollen compartments thus imposes a physical barrier to fusion and thereby accounts for the selective sensitivity of certain viruses to PIKfyve inhibition.

## RESULTS

### Apilimod inhibits infection by VSV-eGFP-SARS-CoV-2 and VSV-MeGFP-ZEBOV but not by VSV-eGFP-G

We first confirmed that infection by VSV-SARS-CoV-2 or VSV-ZEBOV requires PIKfyve activity, whereas infection by VSV-G does not. We inoculated Vero cells not expressing TMPRSS2 with three recombinant VSVs: (i) VSV-eGFP-G, which expresses soluble eGFP and the parental VSV G protein; (ii) VSV-eGFP–SARS-CoV-2, which expresses soluble eGFP and in which the SARS-CoV-2 spike replaces G; and (iii) VSV-MeGFP–ZEBOV, which expresses eGFP fused to the viral matrix protein (M) and carries the Zaire ebolavirus GP instead of G. All three viruses deliver their genomes into the cytosol via endosomal fusion: for VSV, fusion is directly triggered by low pH, whereas Ebola and SARS-CoV-2 Spike first require their proteolytic activation by endosomal proteases, whose activity also requires low pH (13).

We measured infectivity in Vero cells using a single-cell imaging assay that reports expression of virally encoded cytosolic eGFP or MeGFP (13), following the incubation protocol shown in Fig. 1A. To assess the role of endosomal PIKfyve on infection, we treated the Vero cells with apilimod, a selective inhibitor of PIKfyve lipid kinase activity (22). As shown in Figs. 1B-D, apilimod fully inhibited infection by VSV-eGFP–SARS-CoV-2 and VSV-MeGFP–ZEBOV but had no effect on VSV-eGFP-G.

**Figure 1.**
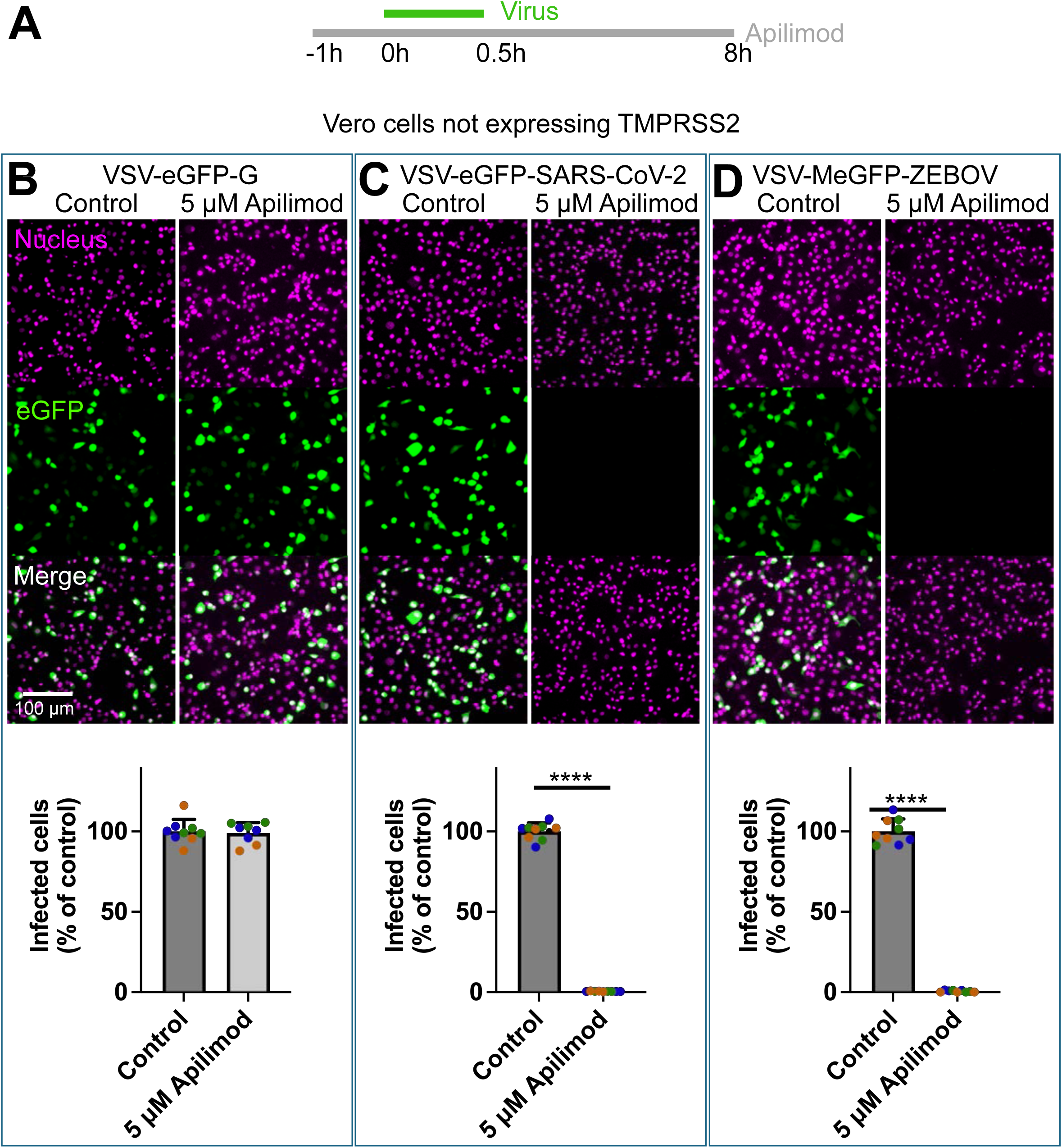
Apilimod inhibits infection by VSV chimeras bearing SARS-CoV-2 or ZEBOV glycoproteins, but not by VSV-G. **(A)** Schematic of the infection assay. Vero cells lacking TMPRSS2 were pretreated with 5 µM Apilimod for 1 h, followed by a 30-min incubation with VSV-eGFP-G, VSV-eGFP–SARS-CoV-2, or VSV-MeGFP–ZEBOV at an approximate MOI of 0.5. in the continued presence of 5 µM apilimod. Unbound virus was removed by three consecutive washes (each containing 5 µM apilimod), and cells were maintained in 5 µM apilimod throughout the experiment. At 7.5 h postinfection, cells were fixed, and infection was quantified as the percentage of eGFP-positive cells relative to total nuclei (Hoechst Janelia 646 stain), using automated image analysis encoded by a script in FIJI. **(B–D)** Representative maximum intensity z-projections (top) show cytosolic eGFP (green) and nuclear Hoechst Janelia 646 (magenta) in cells infected with the indicated chimeras, in the presence or absence of apilimod. Quantification (bottom) shows the percentage of infected cells normalized to the untreated control for each biological replicate (n = 3; shown in distinct colors). Technical triplicates appear as dots of the same color. Approximately 16,000 cells were analyzed per condition. Bars represent mean ± SD. Statistical significance was assessed by unpaired t test; ****P < 0.0001. Comparisons with P > 0.05 are not shown.

### Compartment-specific endosomal swelling

Apilimod treatment of SVG-A cells co-expressing gene-edited fluorescent chimeras of mScarlet-EEA1 (a marker of early endosomes) and NPC1-Halo (a marker of late endosomes and lysosomes) causes swelling of the latter compartment(s), but not of the former (13). We confirmed here that we can detect this compartment-specific swelling in SVG-A cells by lattice light-sheet microscopy (LLSM) (Fig. S1). After apilimod treatment, endosomes labeled with mScarlet alone remained as small, diffraction limited puncta, while those labeled with NPC1-Halo, as well as those containing both labels, became substantially enlarged.

We examined dose dependence of enlargement as follows. We loaded cells with Alexa Fluor 568-labeled dextran (Dextran-AF568), which accumulates in late endosomes and lysosomes, and measured the radii of individual endolysosomes across a range of apilimod concentrations (Fig. 2). In untreated cells, late endosomes and lysosomes had a narrow distribution of radii centered at ∼0.5μm (range 0.3-1.0μm: Fig. 2 B, C). Apilimod induced a concentration-dependent increase in radius, yielding a bimodal distribution with peaks centered at ∼0.75μm and at ∼1.25μm. Applying a 90th percentile cutoff (0.725μm) to score a position as “swollen” yielded the dose-dependent frequency of enlargement shown in Fig. 2 C, D.

**Figure 2.**
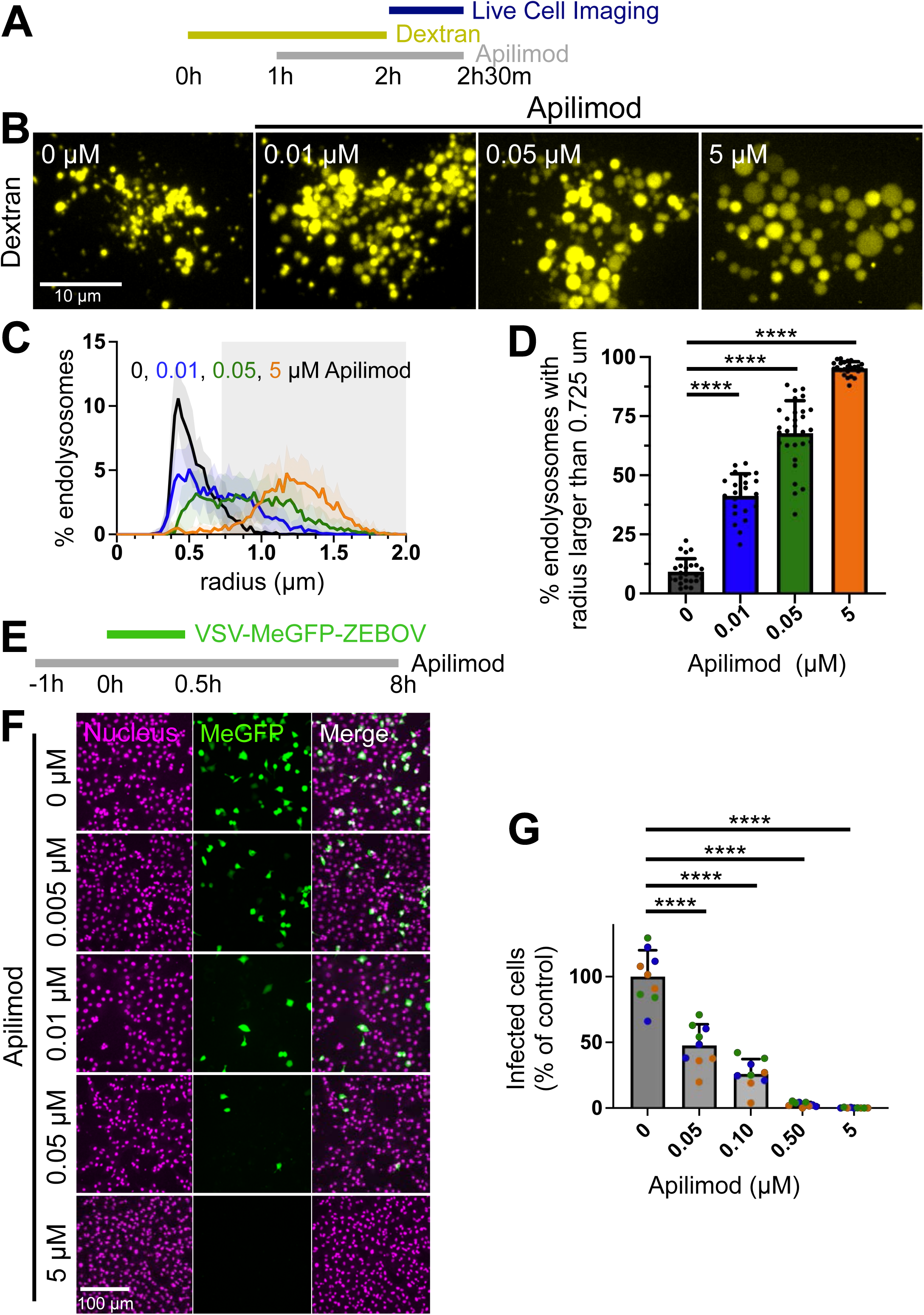
Apilimod induces dose-dependent endolysosome enlargement and inhibits VSV-MeGFP–ZEBOV infection. **(A)** Schematic of the endolysosome enlargement assay. Vero cells not expressing TMPRSS2 were incubated with 10 µM Alexa Fluor 568–labeled dextran (Dextran-AF568) for 2 h, followed by washes to remove unbound dextran. Cells were then treated with apilimod at the indicated concentrations for 1 h and imaged live during a 30 min period, also in the presence of apilimod, using 3D spinning disk confocal microscopy. Dextran-AF568 was used as a fluid-phase marker of endolysosomes. **(B)** Representative maximum intensity z-projection images showing Dextran-AF568 signal (yellow) in cells treated with 0, 0.01, 0.05, or 5 µM apilimod. Enlarged endolysosomes are evident at higher concentrations. Scale bar, 10 µm. **(C)** Distribution of endolysosome radii in cells treated with increasing concentrations of apilimod. Solid lines represent the mean across all cells per condition; shaded regions denote SD. Cell counts: N = 24 (0 µM), 22 (0.01 µM), 29 (0.05 µM), and 24 (5 µM). The gray shaded rectangular region marks the 90th percentile of endolysosome radii with a radius >0.725 µm in the control group (0 µM apilimod). **(D)** Percentage of endolysosomes with a radius > 0.725 µm, plotted against apilimod concentration. Bars represent mean ± SD. Statistical significance was determined by one-way ANOVA compared to control; ****P < 0.0001. **(E)** Schematic of the infection assay. Vero cells not expressing TMPRSS2 were pretreated with 0, 0.005, 0.01, 0.05, or 5 µM apilimod for 1 h, followed by a 30-min infection with VSV-MeGFP–ZEBOV (MOI ∼ 0.5) in the continued presence of apilimod. Unbound virus was removed by three washes (each containing apilimod), and cells were maintained in drug for the remainder of the experiment. At 7.5 h postinfection, cells were fixed, and fraction of infected cell was determined by establishing the of cells expressing MeGFP cytosolic fluorescence with respect to the total number of cells identified by nuclear staining (Hoechst Janelia 646). Automated analysis was performed using FIJI. **(F)** Representative maximum intensity z-projections of images obtained 3D spinning disc confocal fluorescence microscopy showing infected cells (MeGFP, green) and nuclei stained by Hoechst Janelia 646 (magenta) for each apilimod concentration. Scale bar, 100 µm. **(G)** Quantification of infection, plotted as the percentage of infected cells normalized to the untreated control. Each biological replicate (n = 3) is shown in a different color; technical triplicates are represented as dots of the same color. Approximately 16,000 cells were analyzed per condition. Bars represent mean ± SD. Statistical significance was determined by one-way ANOVA; ****P < 0.0001.

### Apilimod-induced late-endosomal swelling inhibits entry of VSV-MeGFP-ZEBOV and VSV-eGFP-SARS-CoV-2 but not of VSV itself

We verified that incoming VSV-MeGFP-ZEBOV particles reach NPC-1 compartments under our experimental conditions by using LLSM with genome-edited SVG-A cells expressing NPC1-Halo (13) labelled with Janelia Fluor 646 HaloTag ligand. Cells were exposed to virions for 10 min, washed, and imaged 40 min or 4 hrs. later at 37 °C, according to the protocol in Fig. 3A. At a multiplicity of infection (MOI) of ∼0.5, each cell imaged at 40 min post inoculation contained 25–30 fluorescent virions in NPC1-positive compartments (Fig. 3B, D). Productive genome entry, monitored by the appearance of MeGFP fluorescence at the nuclear margin (13), was apparent 4 h post-inoculation (Fig. 3E) but not 50 min post-inoculation (Fig. 3B). These observations are consistent with the expectation that ZEBOV GP fuses from late endosomes (23, 24).

**Figure 3.**
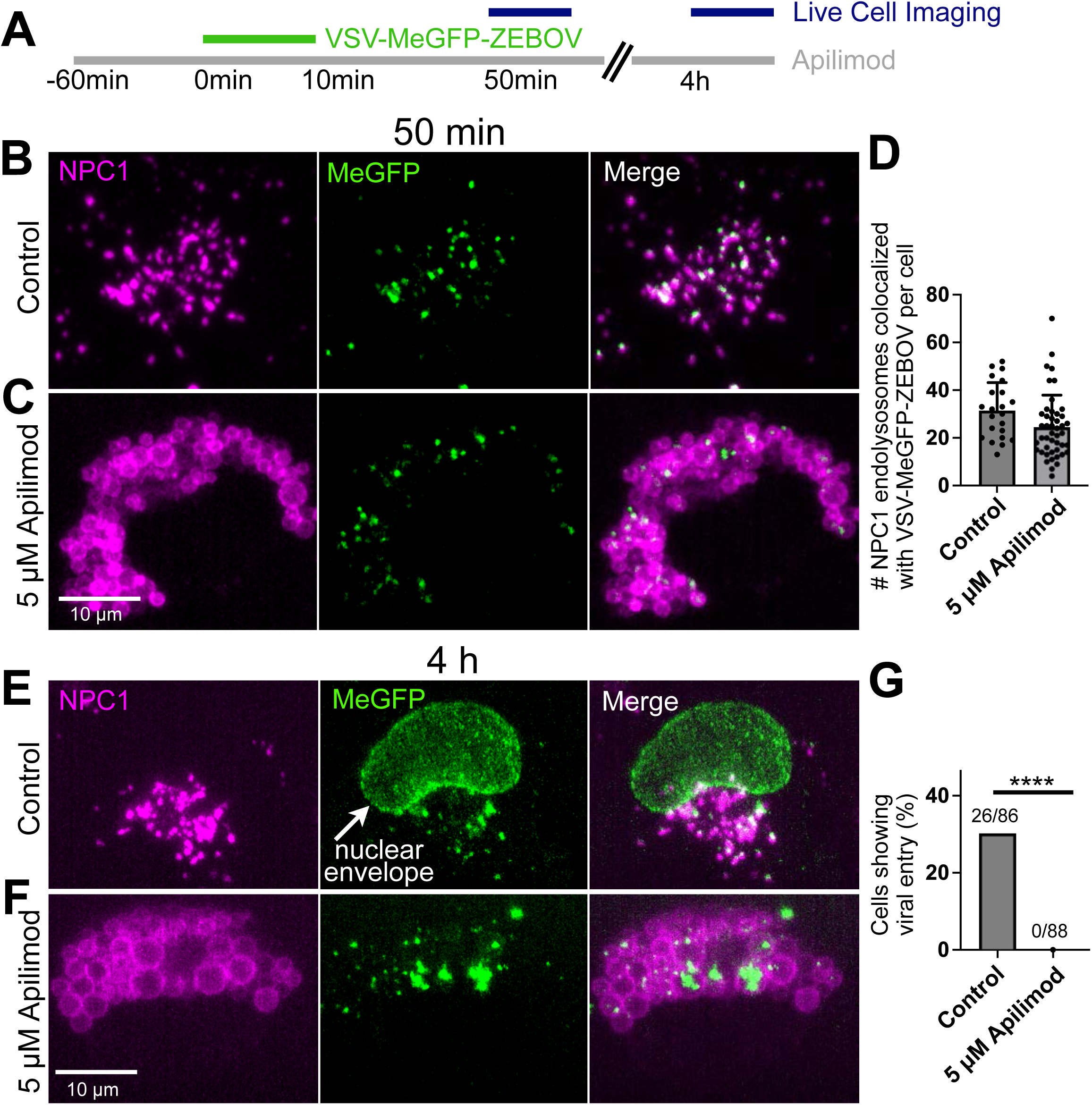
Apilimod does not impair VSV-MeGFP–ZEBOV trafficking to NPC1 compartments but blocks cytosolic escape and infection. **(A)** Schematic of the infection assay. Gene-edited SVG-A cells expressing NPC1-Halo were pretreated with 5 µM apilimod for 1 h, followed by a 10-min incubation with VSV-MeGFP–ZEBOV (MOI ∼0.5) in the continued presence of 5 µM apilimod. Unbound virus was removed by three washes (each containing 5 µM apilimod), and cells were maintained in drug for the remainder of the experiment. Live-cell imaging using lattice light-sheet microscopy was performed at 50 min and 4 h postinfection. Volumetric time-lapse series were acquired every 4 s for 10 min. **(B, C)** Representative maximum intensity z-projections from the first frame of volumetric time series acquired at 50 min postinfection in **(B)** absence (control) or presence **(C)** of 5 µM apilimod. NPC1-Halo labeled with JFX 646 is shown in magenta, and VSV-MeGFP–ZEBOV in green. Scale bar, 10 µm. **(D)** Quantification of NPC1-positive compartments colocalizing with VSV-MeGFP–ZEBOV at 50 min postinfection. Each point represents a single cell (n = 22 control; n = 46 apilimod-treated). Statistical significance was assessed using an unpaired t test; P > 0.05 was considered not significant and is not shown. **(E, F)** Representative maximum intensity z-projections from 4 h postinfection in **(E)** absence (control) or presence **(F)** of 5 µM apilimod. NPC1-Halo labeled with JF 646 is shown in magenta, and VSV-MeGFP–ZEBOV in green. Scale bar, 10 µm. **(G)** Quantification of number of cells showing viral entry based on MeGFP signal at the nuclear margin as in panel **(E).** Statistical significance was assessed by unpaired t test; ****P < 0.0001.

In the presence of 5 μM apilimod, virions trafficked efficiently to NPC1-positive compartments within 50 minutes (Fig. 3C, D), despite pronounced late endosome and lysosome enlargement, showing that PIKfyve inhibition had not disrupted traffic from the cell surface. At 4 h post-inoculation, however, the apilimod-treated cells showed no MeGFP signal at the nuclear periphery (Fig. 3F) and failed to support infection (Fig. 3G). That is, PIKfyve inhibition blocked viral genome release from a (swollen) late endosomal compartment, in agreement with our published observations (13). Infection efficiency decreased with increasing apilimod concentration and endosomal swelling (Fig. 2E, G), further supporting the conclusion that late endosomal enlargement impairs fusion and genome release.

VSV–SARS-CoV-2 entry can proceed through either a full low-pH and cathepsin-dependent stage, in cells lacking TMPRSS2, or a pathway requiring activation by TMPRSS2 on the cell surface (25–27). SARS-CoV-2 entry is apilimod sensitive in cells that lack TMPRSS2 (Fig. 1C), but only partially so in cells that express the protease (28) (Fig. 4, 0% hypo). Thus, when TMPRSS2 is present, the virus appears to follow both entry pathways, and fusion can occur either at the plasma membrane or in early endosomes, after TMPRSS2 cleavage, or alternatively in acidic late endosomes, following cathepsin activation. In its absence, entry is restricted to the late endosomal/lysosomal, cathepsin-dependent route.

**Figure 4.**
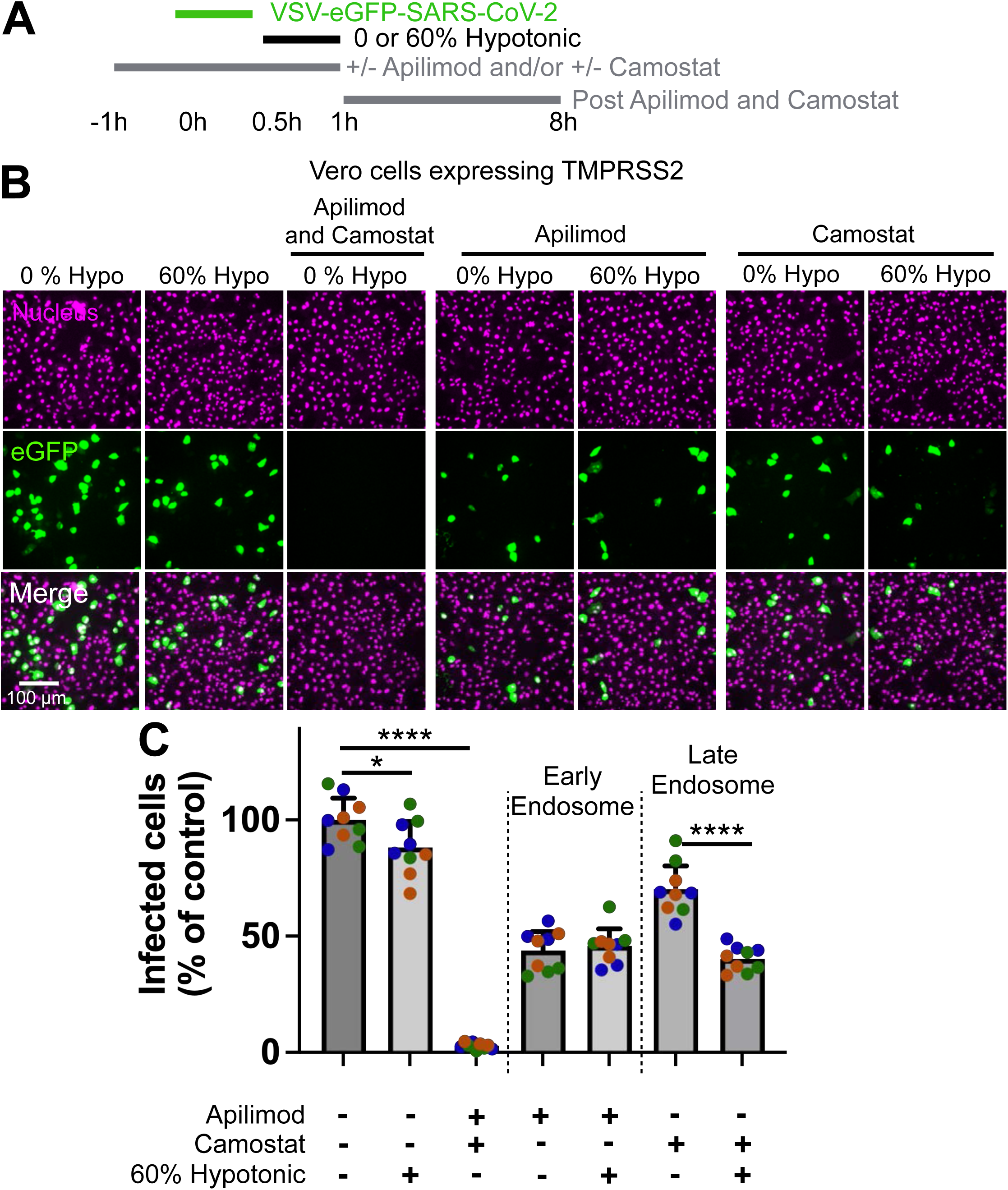
Hypotonic stress partially inhibits VSV-eGFP–SARS-CoV-2 infection via the late endosomal pathway but not early endosome–dependent entry. **(A)** Schematic of the infection assay. Vero cells stably expressing TMPRSS2 were pretreated for 1 h with 5 µM apilimod, 5 µM camostat, both, or DMSO vehicle. Cells were then incubated with VSV-eGFP–SARS-CoV-2 for 30 min (MOI ∼ 0.5) in the continued presence of the same treatment solution. Unbound virus was removed by three washes, each containing the same treatment. Cells were subsequently exposed to 0% or 60% hypotonic medium for 30 min. After treatment, 5 µM apilimod and 5 µM camostat were maintained in the culture medium for the remainder of the experiment (Post apilimod +/- camostat). At 7.5 h postinfection, cells were fixed, and infection was assessed by quantifying eGFP-positive cells relative to total nuclei (Hoechst Janelia 646 stain). Automated analysis was performed using FIJI. **(B)** Representative maximum intensity z-projections show cytosolic eGFP (green) and nuclear Hoechst Janelia 646 (magenta) under the indicated treatment conditions with apilimod or camostat during virion incubation and hypotonic treatment. **(C)** Quantification of infection levels. Data are shown as the percentage of infected cells normalized to untreated control for each biological replicate (n = 3; shown in distinct colors). Technical triplicates are represented as same-colored dots. Approximately 16,000 cells were analyzed per condition. Bars represent mean ± SD. Statistical significance was determined using one-way ANOVA; *P < 0.05, ****P < 0.0001. Comparisons with P > 0.05 are not shown.

We confirmed that apilimod does not inhibit infection by VSV–eGFP–G (Fig.1B) and correlated its resistance with entry from early endosomes. We monitored fusion and cytosolic release, using VSV–PeGFP–G. Expression of eGFP fused to the viral phosphoprotein (P) allowed us to discriminate between fluorescent virions within NPC1-tagged endosomes and those whose genome was released into the cytosol. SVG-A NPC1-Halo–expressing cells were pulse-inoculated and imaged by LLSM at 30- and 40-minutes post-inoculation, with or without 5 μM apilimod (Fig. 5A–C). After 30 min and in both conditions, 8–96 virions per cell had trafficked to NPC1-positive compartments, and ∼20% of P-spots associated with the viral genome had been released into the cytosol (Fig. 5D–F). These results indicate that genome release occurs in early endosomes, with a subset of virions becoming trapped in late endosomes. The similar fraction of cytosolic particles in apilimod-treated and untreated cells is consistent with the resistance of VSV-G–mediated infection to PIKfyve inhibition and agrees with similar results obtained in prior infectivity studies (13).

**Figure 5.**
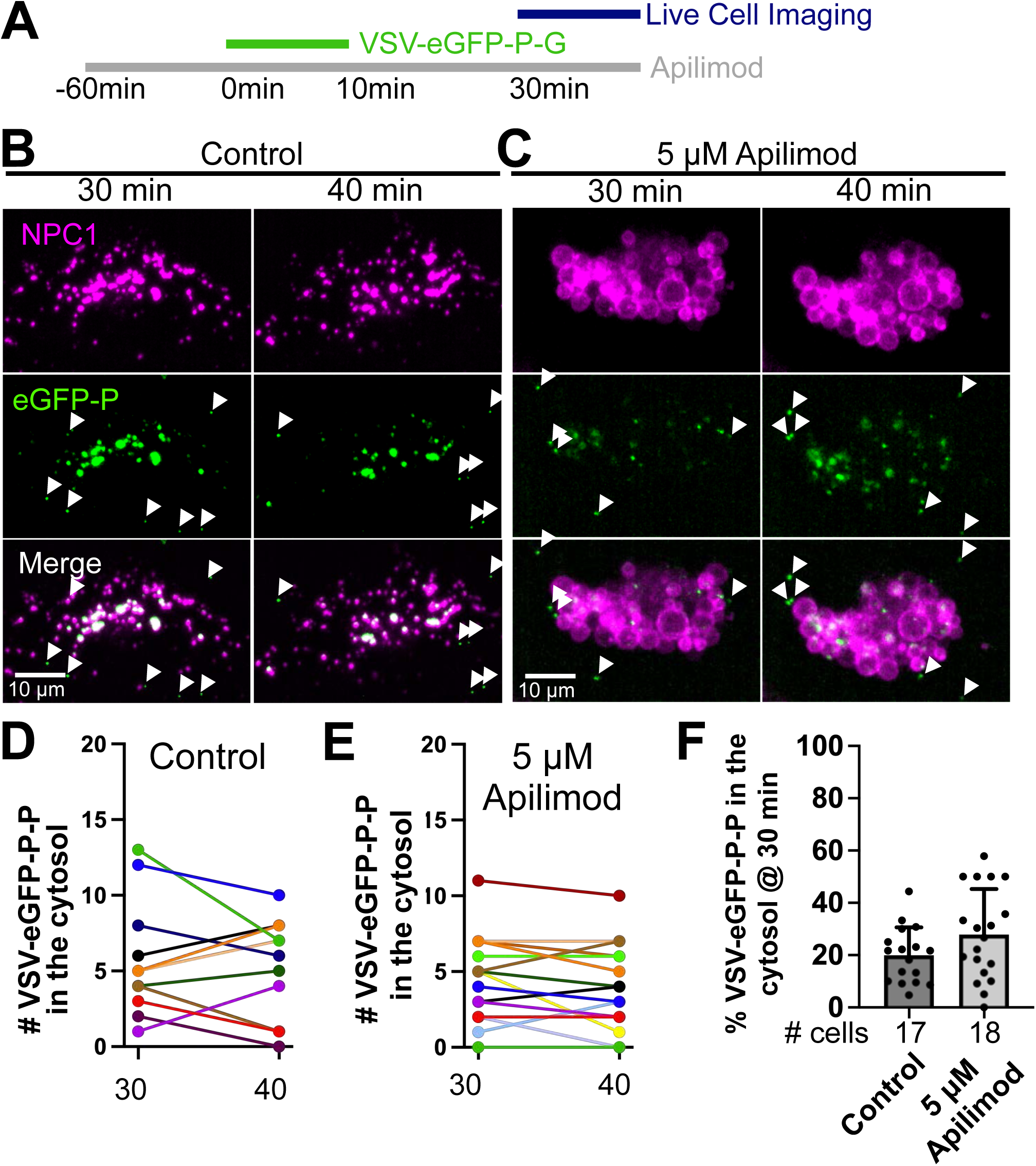
Cytosolic escape and infection by VSV-eGFP–G are not inhibited by apilimod. **(A)** Schematic of the infection assay. Gene-edited SVG-A cells expressing NPC1-Halo were pretreated with 5 µM apilimod for 1 h, then incubated with VSV-eGFP–G (MOI ∼1) for 10 min in the continued presence of drug. Unbound virus was removed by three washes (each containing 5 µM apilimod), and cells were maintained in apilimod for the remainder of the experiment. Live-cell imaging was performed at 30 min postinfection using lattice light-sheet microscopy. Volumetric time-lapse series were acquired every 4 s for 10 min. **(B, C)** Representative maximum intensity projections from the first (30 min) and last (40 min) frames of the time series in the absence **(B)** (control) or presence **(C)** of 5 µM apilimod. NPC1-Halo labeled with JFX 646 is shown in magenta, and VSV-eGFP–G in green. White arrowheads mark virions localized in the cytosol. Scale bar, 10 µm. **(D, E)** Quantification of cytosolic VSV-eGFP–G particles per cell at 30 and 40 min postinfection in **(D)** control and **(E)** apilimod-treated cells. Lines connect values from the same cell at both time points. **(F)** Quantification of the percentage of VSV-eGFP–G particles in the cytosol, with or without apilimod. Statistical significance was assessed by unpaired t test; P > 0.05 was considered not significant and is not shown.

### Endosomal swelling by ammonium accumulation, not PI(3,5)P₂ loss, accounts for blockage of VSV–MeGFP-ZEBOV infection by PIKfyve inhibitors

Under normal conditions, cytosolic glutamine is converted in mitochondria to glutamate and NH₃, which enters endosomes and lysosomes, where acidification by the endo-lysosomal V-ATPase traps it as NH₄⁺ (29). PIKfyve inhibition induces endo-lysosomal swelling through ammonium accumulation and water influx, an effect prevented by glutamine withdrawal from the medium (29) or by inhibition of the V-ATPase (30). We took advantage of this relationship to distinguish between a biochemical consequence of PI(3,5)P₂ depletion by acute apilimod treatment from a broader physical block to infection caused by endosomal swelling, by testing whether the blockade of infection could be reversed by limiting ammonium accumulation.

To validate our experimental system, we imaged SVG-A cells in FluoroBrite medium containing apilimod, with or without supplementation with 4 mM glutamine and FBS. Late endosomes and lysosomes were labeled with internalized Dextran-568 (Fig. 6A). As expected, apilimod-induced endosomal swelling, visually monitored for up to 30 min, was markedly reduced in glutamine-free medium (Fig. 6B). Based on this observation, we inoculated cells for 30 min with VSV-MeGFP-ZEBOV in standard buffer (e.g. containing glutamine), followed by a 30 min incubation with apilimod in the presence or absence of glutamine. Glutamine and 50 nM apilimod were then added for the remainder of the assay to restrict the test of viral entry to the 30 min post-inoculation window, comparing the effect with and without glutamine (Fig. 6C). This step was necessary because prolonged incubation in glutamine-free medium during early infection blocked synthesis of MeGFP encoded by VSV-MeGFP–ZEBOV. VSV-MeGFP–ZEBOV infection was reduced by ∼60% in the presence of both glutamine and apilimod but was minimally affected in glutamine-free medium with apilimod (Fig. 6C–E). These results suggest that inhibition of infection arises from endosomal swelling itself, rather than from loss of PI(3,5)P₂-dependent signaling.

**Figure 6.**
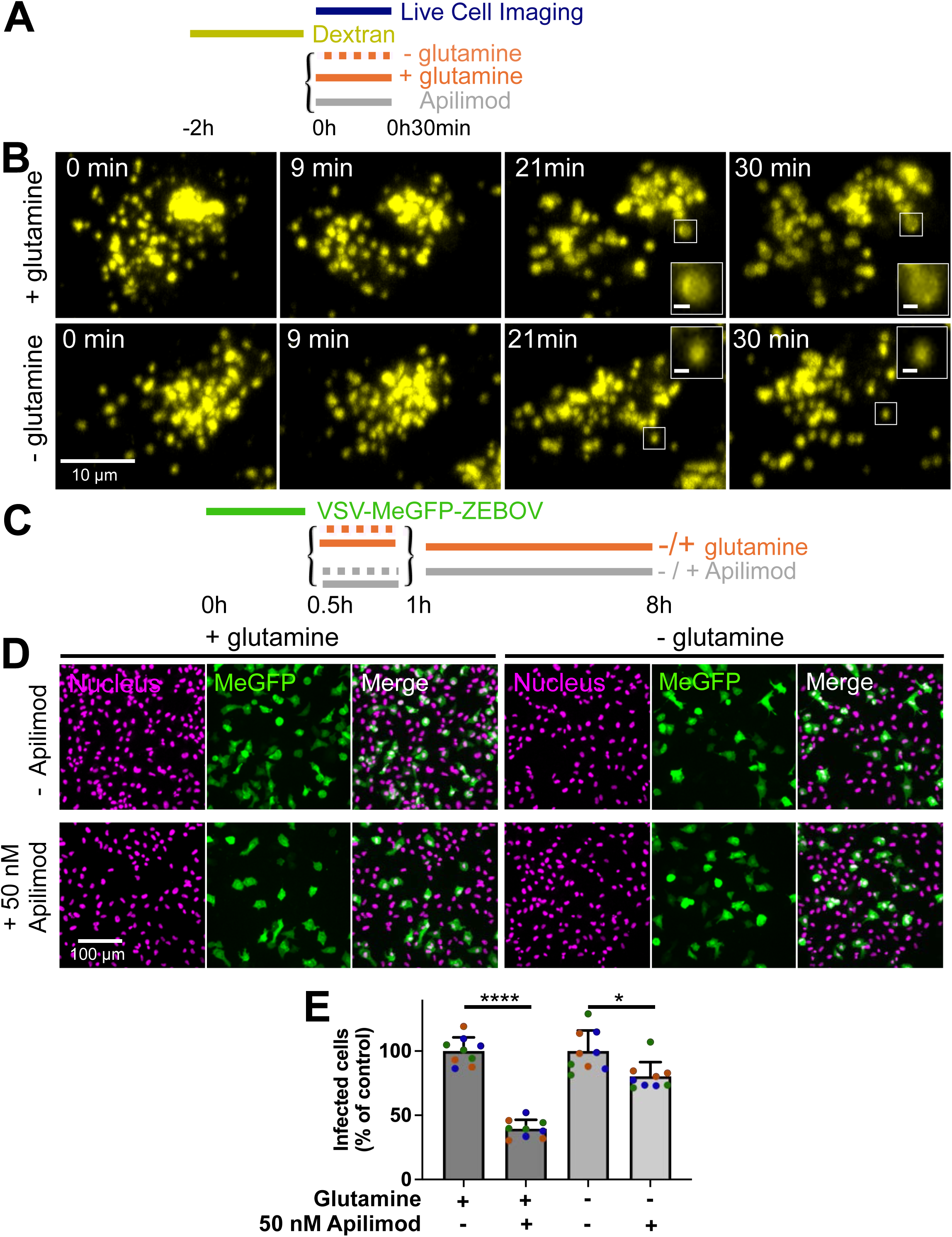
Glutamine depletion moderates apilimod-induced endolysosome enlargement and reduces its antiviral efficacy. **(A)** Schematic of the live-cell assay. Vero cells not expressing TMPRSS2 were incubated with 10 µM Alexa Fluor 568–labeled dextran (Dextran-AF568) for 2 h to label endolysosomes, followed by washes to remove unbound dextran. Cells were then imaged live by lattice light-sheet microscopy during treatment with 50 nM apilimod in either glutamine-containing or glutamine-free medium. Imaging was initiated simultaneously with apilimod addition. **(B)** Representative maximum intensity z-projections from volumetric time series at t = 0, 9, 21, and 30 min post-apilimod addition, with or without glutamine. Dextran-AF568–labeled endolysosomes are shown in yellow. Scale bar, 10 µm and 1 µm in insets. **(C)** Schematic of the infection assay. Vero cells were incubated with VSV-MeGFP–ZEBOV for 30 min, unbound virus was removed by three washes, and cells were treated for 30 min with 50 nM apilimod or DMSO vehicle in either glutamine-containing or glutamine-free medium. Cells were then transferred to glutamine-containing medium with 5 µM apilimod for the remainder of the experiment. At 7.5 h postinfection, cells were fixed, and infection was quantified as the percentage of MeGFP-positive cells relative to total number of cells identified by nuclear staining (Hoechst Janelia 646). Automated analysis was performed using FIJI. **(D)** Representative maximum intensity z-projections showing infected cells (MeGFP, green) and nuclei stained with Hoechst Janelia 646 (magenta) under each condition. **(E)** Quantification of infection. Data are shown as the percentage of infected cells normalized to untreated control. Each biological replicate (n = 3) is shown in a different color; technical triplicates are represented as same-colored dots. Approximately 16,000 cells were analyzed per condition. Bars represent mean ± SD. Statistical significance was assessed by unpaired t test; *P < 0.05, ****P < 0.0001.

### Hypotonic-induced endolysosomal swelling reduces VSV-MeGFP–ZEBOV and VSV-eGFP–SARS-CoV-2 infection

To test directly whether endolysosomal osmotic swelling impairs viral entry, we used a non-pharmacological method to enlarge endosomes and lysosomes during the early phase of infection. Brief incubation of cells in hypotonic medium induces transient swelling of the cytoplasm and intracellular membrane-bound compartments, most prominently late endosomes and lysosomes (31–33). We exploited this osmotic effect to perturb the entry process of VSV-eGFP-G, VSV-MeGFP–ZEBOV and VSV-eGFP–SARS-CoV-2. A single 30-minute hypotonic pulse, initiated 30 minutes post-inoculation, was reversible, did not lead to cell death (Fig. 4B) consistent with prior reports showing no overt toxicity from a similar transient osmotic shock (34), and targeted the entry window while minimizing artifacts from subsequent volume regulation. VSV-eGFP-G infection remained unchanged with or without the treatment (Fig. 4C).

To quantify endosomal swelling, we incubated Vero cells in medium diluted with increasing amounts of water and measured endolysosomal size by spinning-disk confocal imaging of cells preloaded with Alexa Fluor 568–labeled dextran (Dextran-AF568; Fig. 7A). Using the same 90th percentile radius cutoff (0.725 μm) as in the apilimod experiments, we observed a modest increase in enlarged compartments with 30% dilution and a robust increase with 60% dilution, whereas 10% dilution had no detectable effect (Fig. 7B–D).

**Figure 7.**
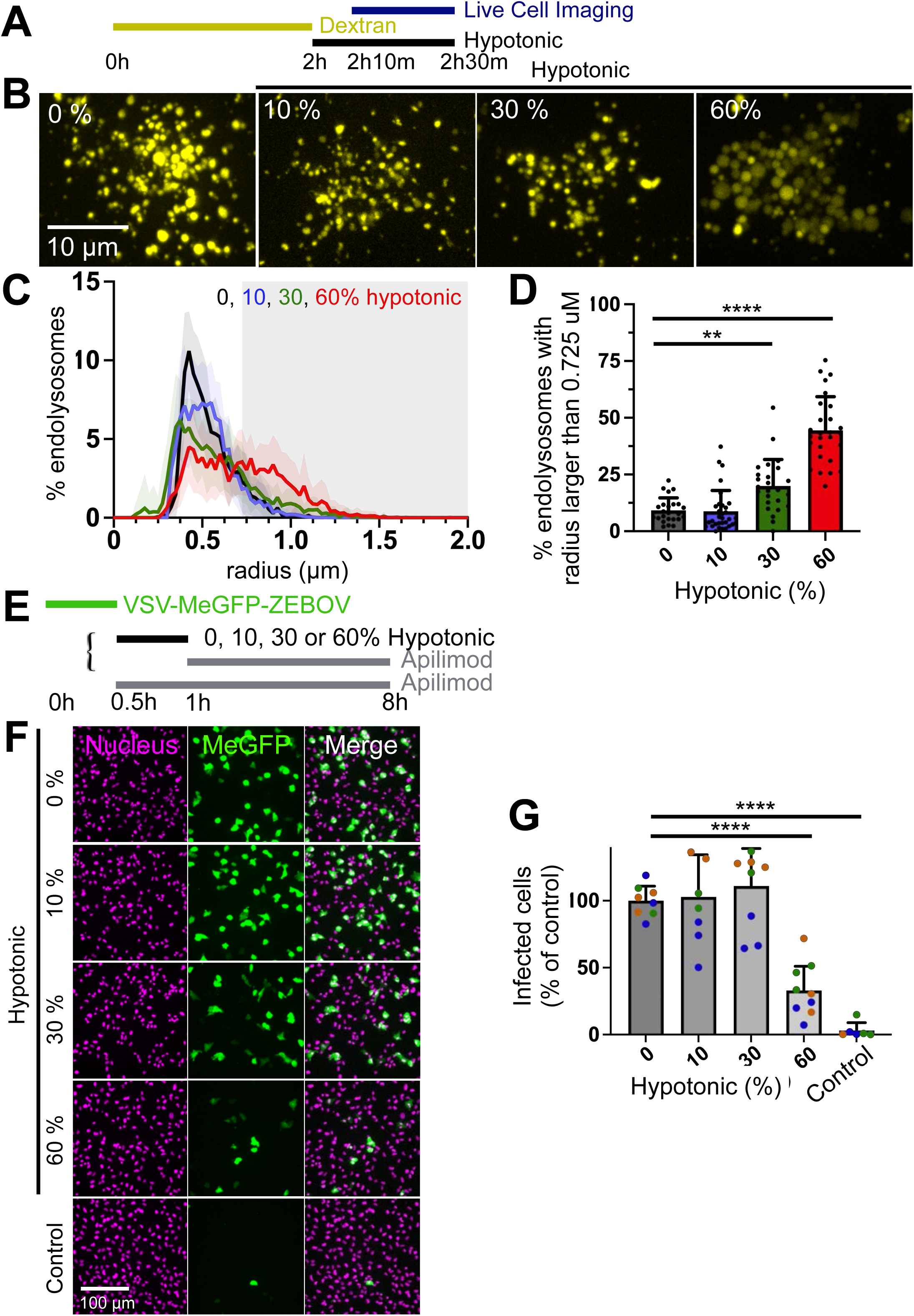
Hypotonic-induced endolysosome enlargement is dose-dependent and correlates with inhibition of VSV-MeGFP–ZEBOV infection. **(A)** Schematic of the endolysosome enlargement assay. Vero cells not expressing TMPRSS2 were incubated with 10 µM Alexa Fluor 568–labeled dextran (Dextran-AF568) for 2 h to label endolysosomes, followed by washes to remove unbound dextran. Cells were then exposed to hypotonic media (0%, 10%, 30%, or 60%) for 10 min prior to imaging, which was performed using live cell spinning disk confocal microscopy. Hypotonic conditions were maintained throughout image acquisition. **(B)** Representative maximum intensity z-projections showing Dextran-AF568 signal (yellow) in cells exposed to increasing hypotonic medium. Enlarged endolysosomes are observed in 30% and 60% conditions. Scale bar, 10 µm. **(C)** Distribution of endolysosome radii for cells treated with 0% (black), 10% (blue), 30% (green), or 60% (red) hypotonic media. Solid lines represent the mean endolysosome radius across all cells per condition; shaded areas denote SD. Cell counts: N = 24 (0%), 32 (10%), 25 (30%), and 27 (60%). The gray shaded region marks the 90th percentile of endolysosome with a radius > 0.725 µm in control cells (0%). **(D)** Percentage of endolysosomes with radius >0.725 µm, plotted against hypotonic media concentration. Bars represent mean ± SD. Statistical significance was assessed by one-way ANOVA compared to control; **P < 0.01, ****P < 0.0001. Comparisons with P > 0.05 are not shown. **(E)** Schematic of the infectivity assay. Vero cells were infected with VSV-MeGFP–ZEBOV (MOI ∼ 0.1) for 30 min. Unbound virus was removed by three washes, and cells were then treated with 0%, 10%, 30%, or 60% hypotonic media or 5 µM apilimod for 30 min. Following treatment, all conditions were maintained in 5 µM apilimod for the duration of the experiment. At 7.5 h postinfection, cells were fixed, and infection was quantified by establishing the ratio of cells expressing MeGFP cytosolic fluorescence with respect to the total number of cells identified by nuclear staining using automated image analysis in FIJI. **(F)** Representative maximum intensity z-projections showing infected cells (MeGFP, green) and nuclei stained with Hoechst Janelia 646 (magenta) for each treatment condition. Scale bar, 100 µm. **(G)** Quantification of infection levels. Data are shown as the percentage of infected cells normalized to the untreated control for each biological replicate (n = 3; distinct colors). Technical triplicates appear as dots of the same color. Approximately 16,000 cells were analyzed per condition. Bars represent mean ± SD. Statistical significance was assessed by one-way ANOVA; ****P < 0.0001. Comparisons with P > 0.05 are not shown.

We next tested the impact of this hypotonic swelling on infection. SVG-A or Vero cells were inoculated with VSV-MeGFP–ZEBOV, VSV-eGFP–SARS-CoV-2, or VSV-eGFP-G, then subjected to 30-minute hypotonic treatments at the indicated dilutions (Figs. 7E-G and 8). To restrict continued viral fusion after the swelling phase, we added Bafilomycin A1 alone (VSV-G), apilimod alone (VSV-MeGFP-ZEBOV) or in combination with apilimod (VSV-SARS-CoV-2) immediately following the hypotonic pulse. Bafilomycin A1 blocks endosomal acidification (35) and thereby prevents low pH-dependent viral membrane fusion. The combination of Bafilomycin A1 and apilimod inhibited fusion of VSV-MeGFP–ZEBOV and VSV-eGFP–SARS-CoV-2. Infection of SVG-A cells by VSV-MeGFP–ZEBOV dropped to ∼40% of control levels following 60% dilution, with no inhibition at lower dilutions (Fig. 7F and G). The same treatment in Vero cells suppressed infection by ∼ 50% for both VSV-eGFP–SARS-CoV-2 and VSV-MeGFP–ZEBOV but had no effect on VSV-eGFP (Fig. 8B-D). The similarity between these results and those obtained with apilimod supports the conclusion that late endosomal swelling mediated by transient hypotonicity rather than inhibition of a specific biochemical pathway selectively blocks infection by viruses that require trafficking to late endosomes, preventing membrane fusion and cytosolic genome delivery.

**Figure 8.**
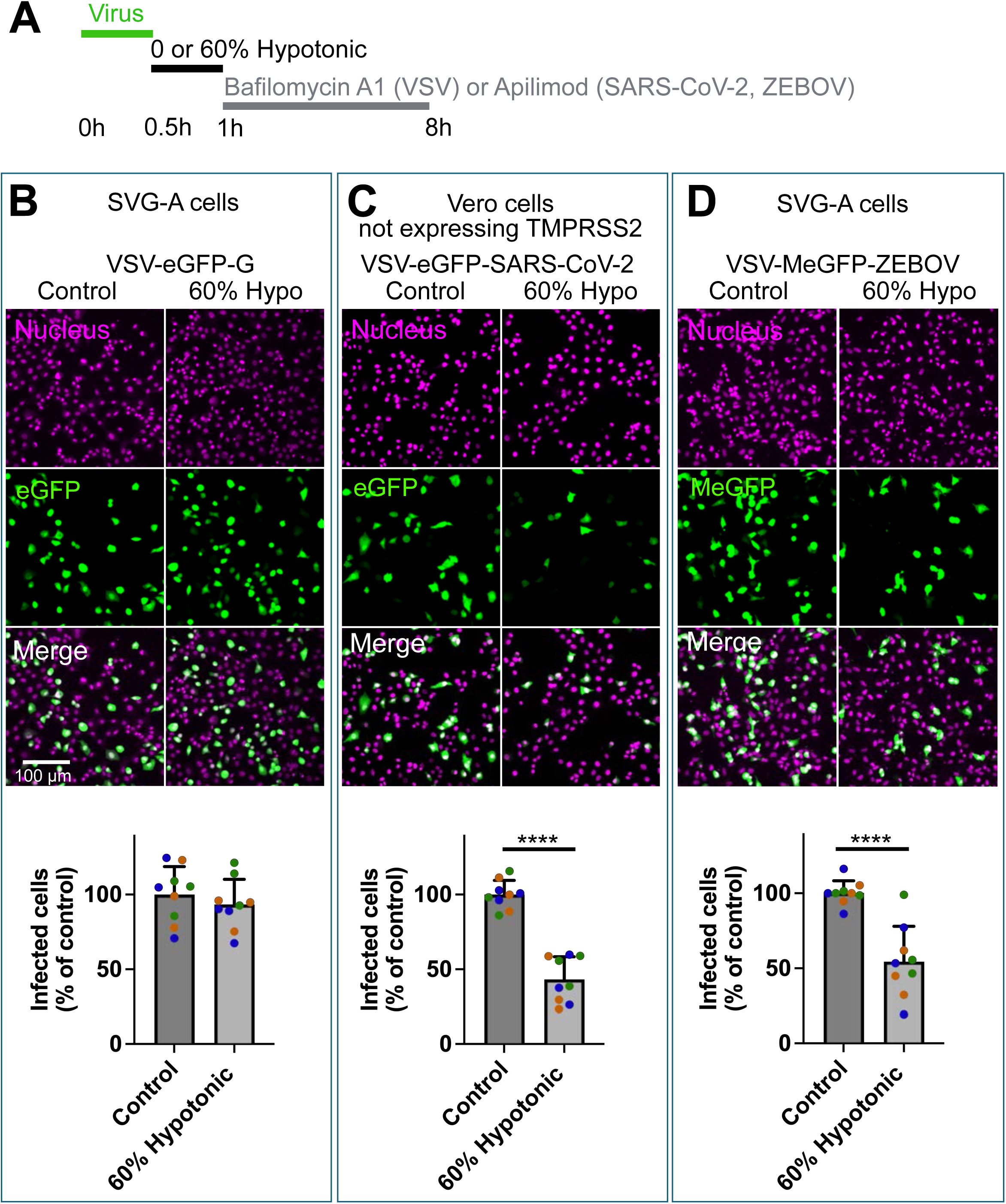
Hypotonic stress partially inhibits infection by VSV-eGFP–SARS-CoV-2 and VSV-MeGFP–ZEBOV, but not by VSV-eGFP–G. **(A)** Schematic of the infection assay. SVG-A or Vero cells not expressing TMPRSS2 were incubated with VSV-eGFP–G, VSV-eGFP–SARS-CoV-2, or VSV-MeGFP–ZEBOV (MOI ∼ 0.5) for 30 min, followed by three washes to remove unbound virus. Cells were then treated with 0% or 60% hypotonic medium for 30 min. Following treatment, cells were maintained with normal medium with either 100 nM Bafilomycin A1 (used only for VSV-eGFP–G samples) or 5 µM apilimod (used for the other virions) for the remainder of the experiment. At 7.5 h postinfection, cells were fixed, and infection was quantified by counting GFP-positive cells relative to total nuclei (Hoechst Janelia 646) using spinning disc confocal fluorescence microscopy. Automated analysis was performed in FIJI. **(B-D)** Representative maximum intensity z-projection images (top) showing infected cells (eGFP, green) and nuclei (magenta) under 0% and 60% hypotonic conditions for each type of virion. Quantification of infection levels. Data (bottom) are shown as the percentage of infected cells normalized to the untreated control for each biological replicate (n = 3; distinct colors). Technical triplicates are represented as same-colored dots. Approximately 16,000 cells were analyzed per condition. Bars represent mean ± SD. Statistical significance was assessed using an unpaired t test; ****P < 0.0001. Comparisons with P > 0.05 are not shown.

To test whether hypotonic treatment inhibits infection primarily through late endosomal swelling, we took advantage of the differential activation of SARS-CoV-2 spike (S): TMPRSS2 activates S at the cell surface/early endosomes, whereas cathepsins do so in late endosomes (25–27). In Vero cells ectopically expressing TMPRSS2, concurrent treatment with camostat (a TMPRSS2 inhibitor that prevents early entry) and apilimod (which blocks late entry) completely inhibited VSV-SARS-CoV-2 infection (Fig. 4A–C) (28). Either compound alone caused the expected partial inhibition (∼60% for apilimod, ∼25% for camostat) consistent with additive effects on distinct entry routes (Fig. 4C) (28).

If hypotonic swelling acts primarily in late endosomes, infection should be strongly inhibited when TMPRSS2 is absent or blocked by camostat, and only weakly inhibited when TMPRSS2 is active. Our results matched these predictions: 60% hypotonic treatment inhibited infection by 50-60% either without TMPRSS2 (Fig. 7D) or with TMPRSS2 plus camostat (Fig. 4C), but only by ∼20% when TMPRSS2 was present without camostat (Fig. 4C). Consistent with late-endosomal fusion, combining hypotonic treatment with apilimod did not further reduce infection (∼60%, Fig. 4C). Conversely, by preventing early entry, camostat sensitized infection to hypotonic swelling (∼20% without vs ∼ 60% with camostat, Fig. 4C). Together, these results support our inference that hypotonic swelling blocks infection by interfering with the late endosomal entry pathway.

### Free and membrane-bound VSV-MeGFP–ZEBOV virions in swollen endosomes

Finally, we assessed whether VSV-MeGFP–ZEBOV virions remain membrane-bound under swelling conditions that block infection using live-cell 3D LLSM imaging of virions within apilimod-induced, enlarged NPC1-Halo containing late endosomes and lysosomes in SVG-A cells (Figs. 3C, F and 9). We did not use hypotonic swelling because both transience and heterogeneity limit resolution and stability during live-cell imaging. In non-swollen endosomes (∼300-500 nm diameter), the point-spread function of the microscope precludes distinguishing virions at the limiting membrane from those in the lumen (Figs. 3B, E). In contrast, apilimod-induced swelling yields larger endosomes of ∼1 μm diameter (Figs. 3C, F), enabling spatial separation of membrane-bound and luminal particles (Figs. 3C, F and 9). Because these compartments remain acidic (12), we expected most virions to remain receptor-bound and fusion-competent.

**Figure 9.**
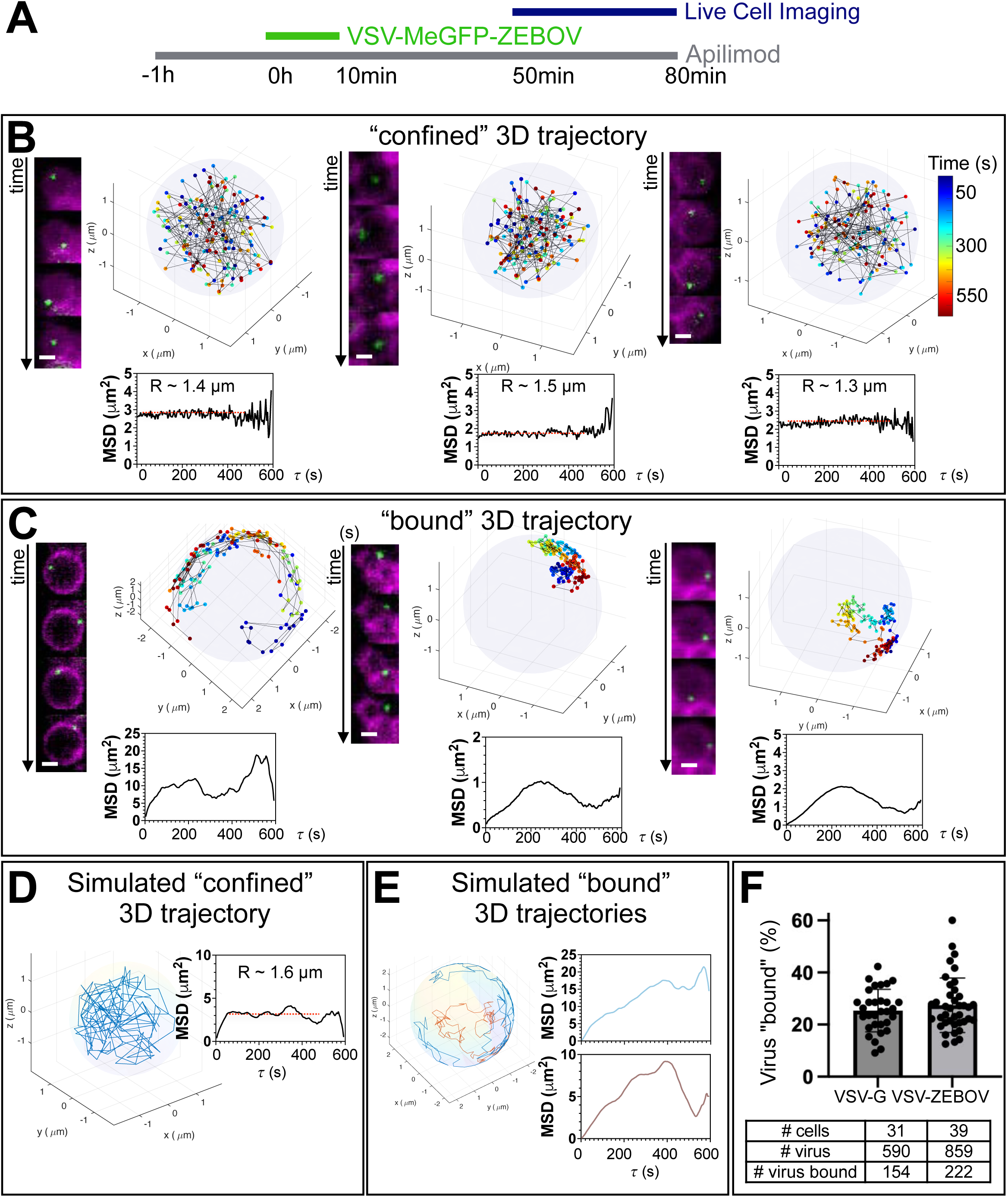
3D tracking of VSV-MeGFP–ZEBOV reveals confined and membrane-bound virion populations within apilimod-induced swollen NPC1-positive endolysosomes. Associated with Movies 1-3. **(A)** Schematic of the infection assay. Gene-edited SVG-A cells expressing NPC1-Halo were pretreated with 5 µM apilimod for 1 h, then incubated with VSV-MeGFP–ZEBOV (10 min, MOI ∼ 1) in the continued presence of 5 µM apilimod. Unbound virus was removed by three washes (each containing 5 µM apilimod), and cells were maintained in apilimod for the duration of the experiment. Live-cell imaging was performed at 50 min postinfection using lattice light-sheet microscopy. 3D time-lapse series were acquired every 4 s over 10 min. **(B, C)** Representative 3D trajectories of individual virions moving within the lumen **(B)** or along the limiting membrane **(C)** of NPC1-positive endolysosomes. NPC1-Halo–positive endolysosomes are shown as light blue circles. Virion positions are color-coded by time (dark blue to dark red). Insets show representative maximum-intensity z-projections from sequential images of an NPC1–Halo–positive endosome labeled with JF646 (magenta) and VSV-MeGFP–ZEBOV (green). Scale bar, 1 um. Mean square displacement (MSD) plots are shown; in **(B)**, the radius of confinement is derived from the MSD plateau using the relationship 6/5 R² (inset). **(D, E)** Simulated 3D trajectories reproduce the observed motion types. **(D)** Confined motion within a spherical volume with step length comparable to the sphere radius. **(E)** Membrane-bound motion on the surface of a rotating sphere at two rotation speeds (blue and brown). Corresponding MSD curves are shown. **(F)** Fraction of VSV-eGFP–G and VSV-MeGFP–ZEBOV virions exhibiting membrane-bound trajectories among all virions entrapped in NPC1-positive endolysosomes. Classification was based on movement patterns in 3D time-lapse maximum-intensity projections.

We used 3D live-cell LLSM to quantify the movement of fluorescent virions inside swollen endolysosomal compartments labeled with endogenously tagged NPC1-Halo in SVG-A cells (Fig. 9). Treatment with 5 μM apilimod produced largely immobile enlarged endosomes, in contrast to the dynamic trafficking observed in untreated cells. Visual inspection of z-projected 3D time series revealed that ∼75% of 859 fluorescent VSV-MeGFP–ZEBOV virions localized to the lumen while the 25% remainder colocalized with the limiting membrane (Fig. 9B, C, F). A similar distribution of virions in the lumen was also observed for VSV-PeGFP-G, which is not sensitive to apilimod treatment (Fig. 9F). In that case, ∼25% of 590 virions remained membrane bound while the rest were in the lumen. These observations were unexpected.

To distinguish luminal from membrane-associated behavior, we tracked the 3D positions of individual VSV-MeGFP–ZEBOV particles over time. Using time-resolved LLSM at 4-second intervals, we followed fluorescent virions within swollen compartments and computed their mean squared displacement (MSD) (representative examples depicted in the insets in Fig. 9). Particles localized to the lumen had nearly constant MSD values (Fig. 9B), consistent with the confinement within a spherical volume, as expected from a plateau value of 6/5 R² (Fig. 9D). In contrast, particles stably associated with the limiting membrane showed constrained motion and MSD profiles consistent with movement of the surface of a rotating sphere (Fig. 9C, E). Variation in particle and compartment dynamics resulted in diverse trajectory patterns. We restricted this analysis to representative examples, as current computational limitations precluded automated processing of larger datasets.

We interpret these single-virion imaging results as follows. In normal acidic endosomes, VSV-eGFP-P-G and VSV-MeGFP–ZEBOV remain stably associated with their respective receptors. VSV-G–mediated fusion occurs primarily in early endosomes, which are not subject to osmotic swelling. Because NPC1 is absent from this compartment, VSV-MeGFP–ZEBOV fusion does not occur there. As VSV-eGFP-P-G and VSV-MeGFP–ZEBOV virions traffic further, arrival in swollen late endosomes correlates with a loss of stable membrane association. We attribute this shift to increased membrane tension, which could impede viral fusion by raising the energetic barrier to the membrane deformations required for hemifusion and full fusion (see Discussion).

In the acidic environment of swollen endosomes, unbound viral glycoproteins in VSV-PeGFP-G or VSV-MeGFP–ZEBOV that have not yet engaged the limiting membrane may undergo a premature low pH–induced conformational change, thus making them unable to rebind and no longer capable of driving fusion. Our data do not imply that swelling intrinsically impairs fusion-peptide exposure or insertion for virions that remain membrane engaged; the “bound” population stays attached to the limiting membrane and can, in principle, insert the fusion peptide upon low-pH activation. Rather, swelling allows us to detect a “loose” luminal virus population whose GP presumably can undergo low-pH triggering without membrane contact, leaving GP in a post-trigger, nonproductive state because the fusion peptide cannot insert into the target membrane in trans. This mechanism would therefore explain both the retention of virions at the membrane and the release of others into the lumen, where they become fusion-incompetent and unable to rebind. For VSV-G, this has little consequence, as fusion typically occurs in the early endosome, e.g. upstream of this compartment. In contrast, for VSV-MeGFP-ZEBOV, the combination of altered membrane mechanics and disrupted NPC1 engagement in late endosomes likely underlies the observed failure to fuse and the consequent block in infection.

## DISCUSSION

We examined why inhibition of the endosomal lipid kinase PIKfyve selectively blocks infection by certain membrane-enveloped viruses, represented here by VSV chimeras bearing the SARS-CoV-2 spike or Ebola virus GP, while leaving parental VSV-G–mediated infection unaffected. Using a combination of quantitative infectivity assays and high-precision, live-cell 3D imaging, we identified a previously unrecognized and mechanistically simple explanation: endosomal membrane swelling acts as a biophysical barrier to viral fusion and genome delivery into the cytosol.

Previous models have attributed the antiviral effects of PIKfyve inhibition primarily to disruptions in endosomal maturation or impaired trafficking of virions to protease-rich, fusion-competent compartments. While consistent with the dependence of SARS-CoV-2 and Ebola virus on late endosomal cathepsin activity and NPC1 function, these models do not account for our earlier observations by live-cell single virion fluorescence microscopy, confirmed here, that these virions trafficked normally to the swollen NPC1-positive compartments under PIKfyve inhibition but failed to initiate productive infection.

Our findings uncouple PI(3,5)P₂ depletion from the block in viral infection. Although apilimod inhibits PIKfyve and reduces PI(3,5)P₂ levels, the antiviral effect against VSV-SARS-CoV-2 and VSV-ZEBOV requires endosomal swelling, a process driven by glutamine-dependent accumulation of NH₄⁺ and consequent osmotic influx. Apilimod treatment in glutamine-depleted media failed to block infection, indicating that PI(3,5)P₂ loss alone is insufficient to impair viral entry. Furthermore, endosomal pH reporters show that apilimod treatment causes at most a modest decrease in endolysosomal luminal pH, arguing that inhibited entry is not due to loss of acidity (36–38). These observations implicate a mechanical disruption of the fusion-permissive environment, rather than defects in endosomal acidification. Thus, the antiviral activity of apilimod depends on a biophysical alteration of endosomal membranes rather than solely on phosphoinositide depletion.

We further demonstrated that transient hypotonic late endosome swelling, in the absence of PIKfyve inhibition, phenocopies the antiviral effect, reinforcing our conclusion that osmotic expansion of endosomal volume rather than disruption of lipid signaling impairs viral entry. These observations together with our observation that apilimod did not hinder the traffic of VSV-G, VSV-ZEBOV or VSV-SARS-CoV2 to the endolysosomal compartment (this study and see also (13)) strengthen the conclusion that the key barrier is not trafficking or maturation failure, but instead a tension-induced mechanical resistance of the endosomal membrane to the deformation required for fusion and pore formation. Fusion requires precise spatial coordination between viral and host membranes, and a membrane environment permissive for local deformation. Swollen endosomes may resist the bending and curvature of the limiting membrane necessary for pore formation, particularly for glycoproteins with stringent conformational checkpoints, such as VSV-G, EBOV-GP, or SARS-CoV-2 spike.

Pore formation during virus–endosome fusion imposes a substantial elastic cost because the membrane must pass through highly curved, nanometer-scale intermediates. In our system, we take the bending/elastic work required to reach a pore-competent state as G_pore_ ≈ 43–65 k_B_*T*. A minimal energetic comparison asks whether the fusion machinery can supply at least this amount. For influenza, used here as the best studied virus example, multiple analyses indicate that productive fusion requires the concerted action of roughly three HA trimers (39–41). A commonly used estimate for the free-energy drop per HA trimer upon completion of the fusogenic refolding (extended intermediate → postfusion hairpin) is ∼34 ± 3 k_B_*T* (42), giving G_HA_(3) ≈ 3 × 34 ≈ 102 k_B_*T* (roughly 92–112 k_B_*T*). On this accounting, three trimers can, in principle, cover the pore-bending requirement and leave a residual margin G_margin_ = G_HA_(3) − G_pore_ ≈ 37 – 59 k_B_*T* to pay other contributions along the pathway. Membrane tension (γ) directly consumes this margin because it adds an area work term to any intermediate that effectively increases membrane area by ΔA, ΔGγ ≈ γΔA.

Two observations argue that the resting limiting membrane of endosomes/lysosomes already operates near the boundary of available slack area. First, because we cannot measure γ directly on small organelles, we use Fluorescence Lifetime Imaging Microscopy (FLIM) values from Lyso Flipper/Flipper-TR targeted to endosomal organelles as an operational reporter of membrane mechanical state (43, 44). Flipper-TR at the cell surface was calibrated in cells by correlating its FLIM values with cell surface membrane tension inferred from tether forces during osmotic perturbations, establishing a monotonic lifetime–tension relationship that approached a high-tension regime as membranes exhaust slack area (44). Lyso Flipper lifetimes of control endolysosomes fall in this high-lifetime regime (43, 45–47), consistent with a “taut” limiting membrane even before swelling. Second, ultrastructure from electron microscopy (9) showed smooth limiting membranes with limited folded/tubular reserve in the endosomes of control cells, again arguing against significant stores of excess area in control conditions and smooth membranes with no folded/tubular membranes during pharmacological inhibition of PIKfyve (vacuolin-1)-mediated distension, also consistent with “taut” membranes.

Thus, acute osmotic swelling should raise membrane tension (γ) because luminal volume increases faster than the limiting membrane can acquire new area. Under constrained area supply, swelling imposes areal strain, ε = ΔA/A_0_, and increases tension approximately as Δγ ≃ K_A_ε, with K_A_ ≈ 250 ± 50 mN·m⁻¹. Even ε = 0.02 yields Δγ ≈ 5 mN·m⁻¹, placing the limiting membrane in a near-maximal-stretch regime. At these tensions, the area work competes directly with the energy margin available from three HA trimers. Requiring G_margin_ ≳ γΔA gives γ_max_ ≈ (37–59) k_B_*T* / ΔA. For a conservative nanometer-scale ΔA ∼ 50–100 nm², γ_max_ ≈ 1.5–4.9 mN·m⁻¹. Thus, if swelling elevates γ to ∼5 mN·m⁻¹, the tension term alone costs ΔG*_γ_* ≈ 60 k_B_*T* for ΔA = 50 nm² (and ≈120 k_B_*T* for ΔA = 100 nm²), exhausting or exceeding the residual budget after paying G_pore_. Under these swollen conditions, three HA trimers cannot supply enough additional work to drive pore formation unless more HA participate, ΔA is substantially smaller, or local tension is transiently relieved.

Although similar constraints on fusion energetics and copy number have been quantified for SNARE-driven fusion (with estimates that roughly 2-3 SNARE complexes are required for content mixing (48)), comparable quantitative bounds are not yet available for the fusion glycoproteins of VSV, SARS-CoV-2, or Ebola virus. We nevertheless infer that the same physical constraint applies for these virions. This tension–energy budget therefore provides a parsimonious mechanistic explanation for why acute endosomal swelling suppresses entry virion entry pathways that normally proceed in relatively non-swollen compartments and thereby prevents infection.

Why is infection by VSV-G unaffected by apilimod treatment or osmotic swelling? The simplest explanation is that VSV-G–mediated fusion and genome release occur in early endosomes, which are not subject to PIKfyve-dependent swelling. Indeed, previous studies using dominant-negative Rab GTPase mutants showed that VSV infection is inhibited by Rab5(S34N), but not by Rab7(T22N), indicating that fusion occurs before endosomal maturation into Rab7-positive compartments (49–51). Inhibition of PIKfyve kinase by apilimod, at the concentrations used here, also had a minimal effect on influenza-virus infection (18), consistent with the effect of dominant-negative Rab5 and Rab7 GTPase mutants indicating that fusion can occur in both early and late endosomes (51); because apilimod does not swell early endosomes, this route remains available for productive fusion. Our proposed mechanism for apilimod resistance is further supported by our finding that apilimod-mediated inhibition of VSV-SARS-CoV-2 infection is only partial in cells expressing TMPRSS2 and only becomes complete in its absence (this study, see also (28)). TMPRSS2-mediated activation of the SARS-CoV-2 spike promotes fusion at the plasma membrane or within early endosomes, bypassing the late endosomal route (25–27). Accordingly, in TMPRSS2-negative cells where both VSV-SARS-CoV-2 and VSV-ZEBOV rely on late endosomal fusion, the extent of inhibition by osmotic challenge is similar. Thus, viral fusion is governed not solely by pH or receptor engagement but also by the trafficking itinerary of the virion and the mechanical state of the endosomal membrane at the site of fusion.

Many enveloped viruses, including filoviruses, coronaviruses, and arenaviruses rely on late endosomal or lysosomal compartments for entry (see review (52)), making them potentially sensitive to perturbations in membrane mechanics. Targeting host pathways that regulate endosomal membrane tension, such as PIKfyve or its lipid effectors, offers a mechanistically distinct antiviral strategy, one that does not interfere with receptor binding or protease activation, but instead acts at the level of fusion competence. By identifying membrane tension as a key regulator of viral entry, our study highlights the utility of integrating biophysical parameters into models of intracellular viral trafficking and fusion.

## MATERIAL AND METHODS

Detailed material and methods are presented in Appendix SI. Briefly, we carried out single-round infection, endolysosome swelling, and virus-tracking experiments using Vero-derived and SVG-A cell lines, including gene-edited SVG-A cells expressing EEA1–mScarlet and NPC1–HaloTag. We used VSV chimeras bearing native VSV-G or substituted glycoproteins (SARS-CoV-2 S or Ebola virus GP) and propagated them in Vero E6 monolayers. For infection assays, we adsorbed virus for 30 min, removed unbound particles by washing, incubated for ∼7–8 h, fixed with paraformaldehyde, and scored infection by eGFP positivity relative to nuclear counts.

To perturb endolysosomal physiology, we inhibited PIKfyve with apilimod in the presence or absence of glutamine or instead imposed transient hypotonic shock; where indicated, we used bafilomycin A1 or camostat. We labeled endolysosomes with internalized fluorescent dextran or NPC1–Halo ligand and quantified compartment size and virus colocalization by 3D fluorescence microscopy (spinning-disk confocal) and lattice light-sheet imaging. We tracked individual virions in 3D, quantified motility by mean square displacement, and used simulations to interpret confinement and endosome-rotation–driven behavior. Image segmentation and quantification used FIJI/LabKit and custom MATLAB workflows; statistics used one-way ANOVA.

### Statistical Analysis

Statistical comparisons between control and experimental groups were performed using one-way ANOVA (parametric). Significance levels are indicated by asterisks: *p < 0.05; **p < 0.01; ***p < 0.001; ****p < 0.0001. Comparisons without annotated p-values were not statistically significant.

## Materials, Data, and Code Availability

Requests for VSV chimeras and their material transfer agreements should be directed to and will be fulfilled by Dr. Sean Whelan. Datasets and custom analysis code are available at https://github.com/kirchhausenlab. For further information, please contact the lead author at.

## ACKNOWLEDGMENTS

We thank S.C. Harrison for editorial help, S. Whelan for support, M. Cornejo Pontelli for generously providing stocks of VSV chimeras used in this study, E. Sitarksa for imaging the samples used for the endolysosome enlargement assay, B. Zuniga for assistance with determination of virus concentration, and members of the Kirchhausen laboratory for support and encouragement. We also acknowledge the L. Lavis lab and the Open the Chemistry team (Janelia Research Campus, HHMI) for the generous gift of Hoechst Janelia 646 and Janelia Fluor 646 (JF646) dyes. The research was supported by a National Institute of General Medical Sciences Maximizing Investigators’ Research Award GM130386 to T. Kirchhausen and AI163019-05 NIH/NIAID Grant Award to S. Whelan and T. Kirchhausen.

The Zeiss Lattice Lightsheet 7 microscope, located in the IID-HSPH BSL-3 Imaging Core established in 2024 at the Harvard Chan School, was acquired with generous funding from the Massachusetts Life Sciences Center to S. Fortune and T. Kirchhausen. Acquisition of the computing hardware including the DGX’s GPU-based computers, CPU clusters, fast access memory, archival servers, and workstations that made possible this study was supported by generous grants from the Massachusetts Life Sciences Center to T. Kirchhausen and by an equipment supplement to the National Institute of General Medical Sciences Maximizing Investigators’ Research Award GM130386 to T. Kirchhausen. Construction of the server room housing the computing hardware was made possible with generous support from the PCMM Program at Boston Children’s Hospital.

**The authors declare no competing financial interests.**

## Author contributions

N. Chow, G. Scanavachi, A. Saminathan and T. Kirchhausen conceptualized and designed the experiments; N. Chow and A. Saminathan carried the cell biological experiments; G. Scanavachi carried the optical image analysis. T. Kirchhausen drafted the manuscript and revised it in close consultation with all co-authors.

## SUPPLEMENT

## Detailed material and methods

### Reagents

The following reagents were used: apilimod (HY-14644, MedChemExpress), Bafilomycin A1 (B1793-2UG, Sigma-Aldrich), Milli-Q water (Z00QSV0US, MilliporeSigma), Minimum Essential Medium (MEM; 10-010-CV, Corning), Dulbecco’s Modified Eagle Medium (DMEM; 10-013-CV, Corning), penicillin-streptomycin (45000-652, VWR), fetal bovine serum (S11150H, Atlanta Biologicals), Alexa Fluor 568 dextran-10K (D-22912, Invitrogen), Janelia Hoechst 646 and Janelia Fluor 646 HaloTag ligand (gifts from Luke Lavis, Janelia Research Campus, HHMI), FluoroBrite DMEM (A1896701, Gibco), HEPES buffer (SH30237.01, Cytiva), and paraformaldehyde (P6148, Sigma-Aldrich).

### Cell Culture

VeroE6-TMPRSS2 cells (a gift from Siyan Ding (1)) and Vero C1008 cells (Vero 76, clone E6; ATCC CRL-1586) were maintained in DMEM supplemented with 10% fetal bovine serum (FBS) and 1% penicillin-streptomycin (P/S) at 37°C in 5% CO₂. SVG-A cells (ATCC CRL-8621) and a homozygous gene-edited derivative expressing EEA1–mScarlet and NPC1–HaloTag from both alleles (2) were cultured in MEM with 10% FBS and 1% P/S under the same conditions. All experiments used cells passaged fewer than 10 times. Cultures were routinely tested and confirmed negative for mycoplasma.

### Production and Propagation of Chimeric Vesicular Stomatitis Viruses

Recombinant vesicular stomatitis viruses (VSVs) encoding fluorescent reporters and distinct glycoproteins were used. VSV-eGFP-G expresses enhanced GFP (eGFP) and retains the native VSV glycoprotein G (3). VSV-eGFP-P-G expresses eGFP fused to the viral phosphoprotein (P) (2), a cofactor that associates with the large (L) polymerase protein within the virion and is required for efficient RNA-dependent RNA synthesis from the ribonucleoprotein (RNP) template. Each virion contains approximately 165 copies of P (4). In VSV-eGFP–SARS-CoV-2, glycoprotein G is replaced by the SARS-CoV-2 spike (S) protein (Wuhan-Hu-1) (5). VSV-MeGFP-ZEBOV encodes the Zaire ebolavirus (ZEBOV) glycoprotein in place of G and expresses eGFP fused to the matrix (M) protein (6). We estimate that each virion contains ∼1200 MeGFP molecules, assuming the number of MeGFP copies approximates that of the M protein in VSV (7).

Chimeric viruses were propagated in confluent Vero E6 monolayers plated in 15 cm dishes. Growth medium (DMEM with 10% FBS) was removed, and cells were rinsed once with DMEM containing 1% P/S to remove residual serum. Cells were infected at a multiplicity of infection (MOI) of ∼0.01 in 10 mL DMEM supplemented with 0.4% FBS and 1% P/S and incubated for 48 h at 37°C, 5% CO₂. Supernatants were collected, clarified by centrifugation at 3580 rpm for 10 min at 4°C (Beckman Coulter GS-6KR), and divided into two portions: one was stored at 4°C for up to one week for titration and infection assays; the other was aliquoted (0.5 mL), flash-frozen in liquid nitrogen, and stored at −80°C for imaging-based experiments.

### Virus infection

Viral supernatant (100 µL) was diluted in serum-free DMEM containing 1% P/S and applied to cells plated in 48-well plates (82051-004, VWR) for 30 min at 37°C. Unbound virus was removed by three washes with DMEM supplemented with 10% FBS and 1% P/S. Cells were then incubated in fresh complete medium for 8 h at 37°C in 5% CO₂. Fixation was performed with 4% paraformaldehyde (PFA) in PBS for 15 min at room temperature. The multiplicity of infection (MOI) was estimated by quantifying eGFP-positive cells relative to the total population (see Infection Analysis). The 16% (w/v) stock PFA solution was prepared by dissolving 16 g of PFA powder (P6148, Sigma-Aldrich) in 80 mL of preheated 1× PBS (60°C) with stirring for 10 min. We added drops of 10 N NaOH (221465, Sigma-Aldrich) every 5 min until the suspension cleared. After cooling to room temperature, we filtered the solution through a grade 1 Whatman™ filter (09-805-1G, Cytiva), adjusted the pH to 7.2 using a calibrated pH meter (Hi 2211, Hanna Instruments), and aliquoted it into 10 mL portions for storage at −20°C.

### Apilimod Treatment and Infection

Vero E6 cells were seeded at 35–40% confluency in sterile 48-well plates (82051-004, VWR) and used the following day. Cells were pretreated for 1 h at 37°C with 0.1% dimethyl sulfoxide (DMSO; vehicle control) or with 0.01, 0.05, or 5 µM apilimod in DMEM supplemented with 10% FBS and 1% P/S. After pretreatment, cells were inoculated with the indicated chimeric virus for 30 min in serum-free DMEM containing 1% P/S and the corresponding concentration of DMSO or apilimod. Unbound virus was removed by three washes with DMEM containing 10% FBS and 1% P/S, followed by incubation for 7 h at 37°C in 5% CO₂. Cells were then washed three times with PBS (0.5 mL) and fixed with 4% PFA in PBS for 15 min at room temperature. Nuclei were labeled by incubation with 100 nM Janelia Hoechst 646 in PBS for 1 h at room temperature. Imaging was performed without washing out the dye.

### Hypotonic treatment and Infection

Hypotonic media were prepared by diluting complete MEM or DMEM with sterile Milli-Q water: 10% (1 mL water + 9 mL medium), 30% (3 mL water + 7 mL medium), and 60% (6 mL water + 4 mL medium).

SVG-A or Vero E6 cells were seeded at 35–40% confluency in 48-well plates and used the following day. Cells were infected VSV-eGFP–SARS-CoV-2 or VSV-MeGFP-ZEBOV at an MOI of ∼0.5 for 30 min in 200 µL of serum-free MEM for SVG-A or DMEM for Vero E6, each supplemented with 1% P/S, at 37°C and 5% CO₂. Unbound virus was removed by three washes with complete medium (MEM or DMEM supplemented with 10% FBS and 1% P/S, 0.5 mL per wash). Infection was then either blocked by adding 100 nM bafilomycin A1 (VSV-eGFP) or 5 µM apilimod (VSV-eGFP–SARS-CoV-2 and VSV-MeGFP-ZEBOV), and allowed to proceed in corresponding serum-free or hypotonic media for 30 min at 37°C. Cells were then returned to complete medium (10% FBS, 1% P/S) and incubated for 7 h at 37°C in 5% CO₂.

Vero cells stably expressing TMPRSS2 were seeded at 35–40% confluency in sterile 48-well plates (82051-004, VWR) and used the next day. Cells were pretreated for 1 h at 37°C in 5% CO₂ with 0.1% DMSO (vehicle) alone or with 5 µM apilimod, 5 µM camostat, or both in DMEM supplemented with 10% FBS and 1% P/S. Following pretreatment, cells were infected with VSV-eGFP-SARS-CoV-2 (MOI ∼0.5) in 200 µL serum-free DMEM (1% P/S) containing the same inhibitor conditions. After 30 min at 37°C, unbound virus was removed by three washes with complete DMEM (10% FBS, 1% P/S; 0.5 mL/wash). Infection was then allowed to proceed for 30 min in 60% hypotonic medium containing 0.1% DMSO alone or with 5 µM apilimod, 5 µM camostat, or both. Cells were subsequently returned to complete DMEM containing the same inhibitor conditions to prevent further infection and incubated for 7 h at 37°C in 5% CO₂.

The experiment ended by washing the cells three times with PBS (0.5 mL), fixing with 4% PFA in PBS for 15 min at room temperature, and staining with 500 nM Janelia Hoechst 646 in PBS for 1 h before imaging.

### Glutamine Deficiency and Infection

SVG-A cells were seeded at 35–40% confluency in sterile 48-well plates (82051-004, VWR) and used the next day. Cells were infected with the VSV-MeGFP-ZEBOV chimeric virus (MOI ∼0.5) for 30 min in serum-free DMEM (including 4 mM glutamine) supplemented with 1% P/S. Unbound virus was removed by three washes with FluoroBrite DMEM supplemented with 10% FBS, 1% P/S (glutamine-free) or DMEM (including 4 mM glutamine) supplemented with 10% FBS and 1% P/S. The cells were incubated in FluoroBrite DMEM (glutamine-free, 10% FBS, 1% P/S) or DMEM (4 mM glutamine,10% FBS, 1% P/S) containing either 0.1% DMSO or 50 nM apilimod for 30 min at 37°C. After three washes with DMEM (including 4 mM glutamine) supplemented with 10% FBS and 1% P/S, cells were incubated for 7 h at 37°C in 5% CO₂, then washed three times with PBS (0.5 mL) and fixed with 4% PFA in PBS for 15 min at room temperature. Nuclei were then labeled by incubating cells with 100 nM Hoechst Janelia 646 in PBS for 1 h at room temperature. Imaging was performed without washing out Hoechst Janelia 646 dye.

### Apilimod, Glutamine, and Hypotonic Treatments and Endolysosomal Size

Vero E6 cells were seeded at 35–40% confluency in 8-well glass-bottom chambers (C8-1.5H-N, Cellvis) and imaged the following day. Endolysosomes were labeled by incubating cells with 10 µM Alexa Fluor 568–dextran in DMEM supplemented with 10% FBS and 1% P/S for 2 h at 37°C in 5% CO₂. Apilimod (final concentration of 0.01, 0.05 or 5 µM in 0.1 % DMSO) was added for 1 h before and then during the Dextran uptake period, after which cells were washed three times with complete DMEM media (supplemented with 10% FBS and 1% P/S) to remove extracellular dye, followed by live cell imaging in FluoroBrite DMEM supplemented with 5% FBS, 1% P/S, and 25 mM HEPES (pH 7.2) including apilimod at the indicated concentrations. Experimental conditions were initiated in 15-minute intervals across the 8-well chamber. Each well was imaged for 10 minutes before applying the same washing and medium exchange protocol to the next well.

To study the effect of glutamine removal on apilimod induced endolysosome expansion, SVG-A cells were seeded at 35-40% confluency in 8-well glass bottom chambers (C8-1.5H-N, Cellvis) for the following day. Endolysosomes were pre-labeled with 10 µM Alexa Fluor 568–dextran in DMEM supplemented with 10% FBS and 1% P/S for 2 h at 37°C in 5% CO₂. Prior imaging, media was replaced with 50 nM of apilimod in FluoroBrite DMEM supplemented with 10% FBS, 1% P/S (glutamine-free) or DMEM supplemented with 10% FBS and 1% P/S, (including 4 mM glutamine). and timelapse images were acquired in lattice light sheet microscopy to record apilimod induced expansion of dextran labeled compartments.

For osmotic expansion assays, we replaced the imaging medium with hypotonic solutions (10%, 30%, or 60%) and immediately initiated live-cell imaging on a spinning disk confocal microscope at 2-minute intervals for 30 minutes.

### Virus Tracking

Gene-edited SVG-A cells expressing EEA1–mScarlet and NPC1–HaloTag chimeras instead of the native proteins were seeded at 35–40% confluency in 8-well glass-bottom chambers and imaged the following day. NPC1 compartments were labeled by incubating cells with 500 nM Janelia Fluor 646 HaloTag ligand (JFX 646) in MEM supplemented with 10% FBS and 1% P/S for 15 min at 37°C, followed by three washes with the same medium.

Cells were then pretreated for 1 h with 0.1% DMSO (vehicle) or 5 µM apilimod in complete MEM. After pretreatment, cells were incubated with the indicated virus in the same medium for 10 min at 37°C, washed three times, and returned to fresh medium containing the same drug conditions for an additional 50 min. Live 3D imaging was performed using a lattice light-sheet microscope in FluoroBrite DMEM supplemented with 5% FBS, 1% P/S, 25 mM HEPES (pH 7.2), 0.1% DMSO, and 5 µM apilimod.

### Fluorescence Microscopy

#### Imaging of Fixed Samples to Assess Primary Infection

Fixed-cell imaging was performed using a Zeiss Axiovert 200M inverted microscope (Carl Zeiss) equipped with a CSU-22 spinning disk confocal unit (Yokogawa) and controlled by the Marianas imaging platform (3i, Intelligent Imaging Innovations). A 10× objective with a 2× magnifier was used. Samples were excited with 491 nm and 660 nm solid-state lasers (50 mW each), and fluorescence was detected using 525/50 and 680LP emission filters (Semrock), respectively.

Z-stacks (12 optical sections at 3 µm intervals) were acquired with a piezoelectric Z-stage (PZ-2000) and encoded XY stage (MS-2000, Applied Scientific Instruments). Images were captured at 50% laser power (0.68 mW at the back aperture) with 200 ms exposure per plane using a QuantEM 512SC air-cooled CCD camera (Photometrics).

#### Live-Cell 3D Imaging of Endosomes and Lysosomes

Live imaging was performed using a Zeiss Axio Observer Z1 inverted microscope equipped with environmental control (37°C, 5% CO₂) and a CSU-X1 spinning disk confocal unit (Yokogawa), operated via the Marianas (3i) platform. The system included a spherical aberration correction module and a 3i LaserStack with diode lasers at 488, 560 (150 mW), and 640 nm (100 mW). Emission was filtered using 525/40, 609/54, and 692/40 filters (Semrock).

Z-stacks comprising 75 optical planes at 0.270 µm spacing were acquired with 20 ms exposure per plane using a dual-camera sCMOS system (Prime 95B, Teledyne Photometrics). A custom bandpass dichroic (Semrock; transmit 350–560 nm and 641–800 nm, reflect 561–640 nm) enabled simultaneous multichannel imaging. Final XY resolution was 0.145 × 0.145 µm/pixel.

#### Live-Cell 3D Lattice Light Sheet Microscopy for Virus Tracking

Lattice light sheet imaging was performed using a Zeiss Lattice Lightsheet 7 system with a dithered lattice beam (30 µm × 0.7 µm). Cells were maintained at 37°C in a humidified chamber with 5% CO₂. Illumination was provided by 488 nm (10 mW, 2 mW at pupil), 561 nm (10 mW, 2 mW), or 640 nm (5 mW, 1 mW) lasers, typically operated at 1–1.5% power.

A 13.3×, NA 0.4 objective (Zeiss) was used for illumination, and a 44.83×, NA 1.0 objective (Zeiss) for detection. Z-stacks were acquired using sample scanning with 0.25 µm optical sectioning and 0.5 µm axial spacing. Each plane was captured with a 5 ms exposure using dual ORCA-Fusion sCMOS cameras (Hamamatsu), yielding a final XY resolution of 0.145 × 0.145 µm/pixel.

Time-lapse imaging was performed at 3 s intervals for virus tracking and at 3 min intervals for apilimod-induced endosome and lysosome expansion, over durations of 5–45 min. Raw image stacks were deskewed using ZEISS ZEN 3.11 software prior to analysis.

#### Image Analysis

##### Infectivity assay

Z-stacks (12 optical sections) were acquired at 20× magnification to capture cytosolic eGFP signal from infected cells and nuclear staining with Hoechst 646, imaged 8 h post-inoculation (see Fluorescence Microscopy). This protocol ensured that the signal reflected a single round of infection (2). Maximum intensity projections were generated using FIJI. Nuclear segmentation was performed using the LabKit plugin in FIJI (8) to determine the number of cells per field and to generate nuclear masks for infection scoring.

A cell was classified as infected if the eGFP signal within the nuclear mask (z-maximum projection) exceeded a threshold defined by the signal in the eGFP channel from uninfected control samples. The percentage of infected cells was calculated as the number of eGFP-positive nuclei relative to the total number of segmented nuclei, using a custom FIJI script. Results were validated by visual inspection of a representative subset. A minimum of 16,000 cells per condition were analyzed, derived from three independent experiments, each performed in triplicate.

#### Colocalization Between NPC1–Halo–Labeled Compartments and Fluorescently Tagged Virions

Z-stacks from single-timepoint live-cell 3D lattice light sheet images of cells inoculated with M–eGFP–tagged virions were used to assess colocalization with NPC1–Halo–labeled endolysosomal compartments. Virus particles, detected as diffraction-limited spots, were identified using the MATLAB-based 3D CME Analysis software (9). For each virion, mean fluorescence intensity in the NPC1–Halo channel was measured in a 3 × 3 × 3 voxel cube centered on the particle. A virion was classified as colocalized if the NPC1–Halo signal within this volume exceeded the background signal measured in a cytosolic region devoid of NPC1–Halo signal. Analysis was automated using a custom MATLAB script and manually validated in FIJI.

#### Measurement of Endosome and Lysosome Radii

Endosomes and lysosomes labeled with Alexa Fluor 568–dextran internalized for 2 hr were imaged volumetrically using 3D live-cell spinning disk confocal microscopy. Centroid positions of labeled compartments were identified using the 3D CME Analysis software (9). At the z-plane corresponding to each centroid, radial intensity profiles were calculated in the XY plane by measuring the distance from the centroid to the point at which fluorescence intensity declined to 30% of the peak value. Measurements were taken in 2° increments over 360° (180 measurements per compartment), and the mean radial distance was reported as the organelle radius.

#### 3D Tracking of Virions Within NPC1–Halo–Labeled Endolysosomes

Live-cell 3D time-lapse datasets acquired using lattice light sheet microscopy were used to track the motion of M–eGFP–tagged virions within apilimod-induced swollen endolysosomes labeled with NPC1–Halo. The centroid and fluorescence intensity of each diffraction-limited virion were determined at each time point using the 3D CME Analysis software (9). Virion trajectories were assembled by linking detections across frames based on two criteria:

i. The virion remained within a 3D spatial window centered on an NPC1–Halo–positive endolysosome, defined in XY as a 25 × 25-pixel region (∼2.5 × 2.5 µm), centered on the centroid position of the compartment identified using the Multi-Template Matching plugin in FIJI (10). The Z-position of the endolysosome centroid was assigned as the plane of maximal NPC1–Halo signal within ±4 z-planes.
ii. The virion’s fluorescence intensity varied by no more than ± 30% between consecutive time points.

We computed the mean square displacement (MSD) from time series of 3D centroid positions for each virion associated with an enlarged endosome. For each trajectory, we calculated the time-averaged squared displacement as a function of lag time. We classified motility by the lag-time dependence of the MSD. Confined motion showed a brief initial rise followed by a plateau, as confirmed by simulation. In contrast, virions bound to structural elements within the enlarged endosome (e.g., the limiting membrane) lacked a plateau and, after the initial rise, exhibited broad variation in MSD with lag time. Simulations show that rotation of the enlarged endosome accounts for this broad variation.

Confinement of a trajectory within a given endolysosome was validated by visual inspection in FIJI, using maximum intensity projections overlaid with bounding boxes centered on the XY coordinates of the NPC1–Halo signal.

**Fig. S1.**
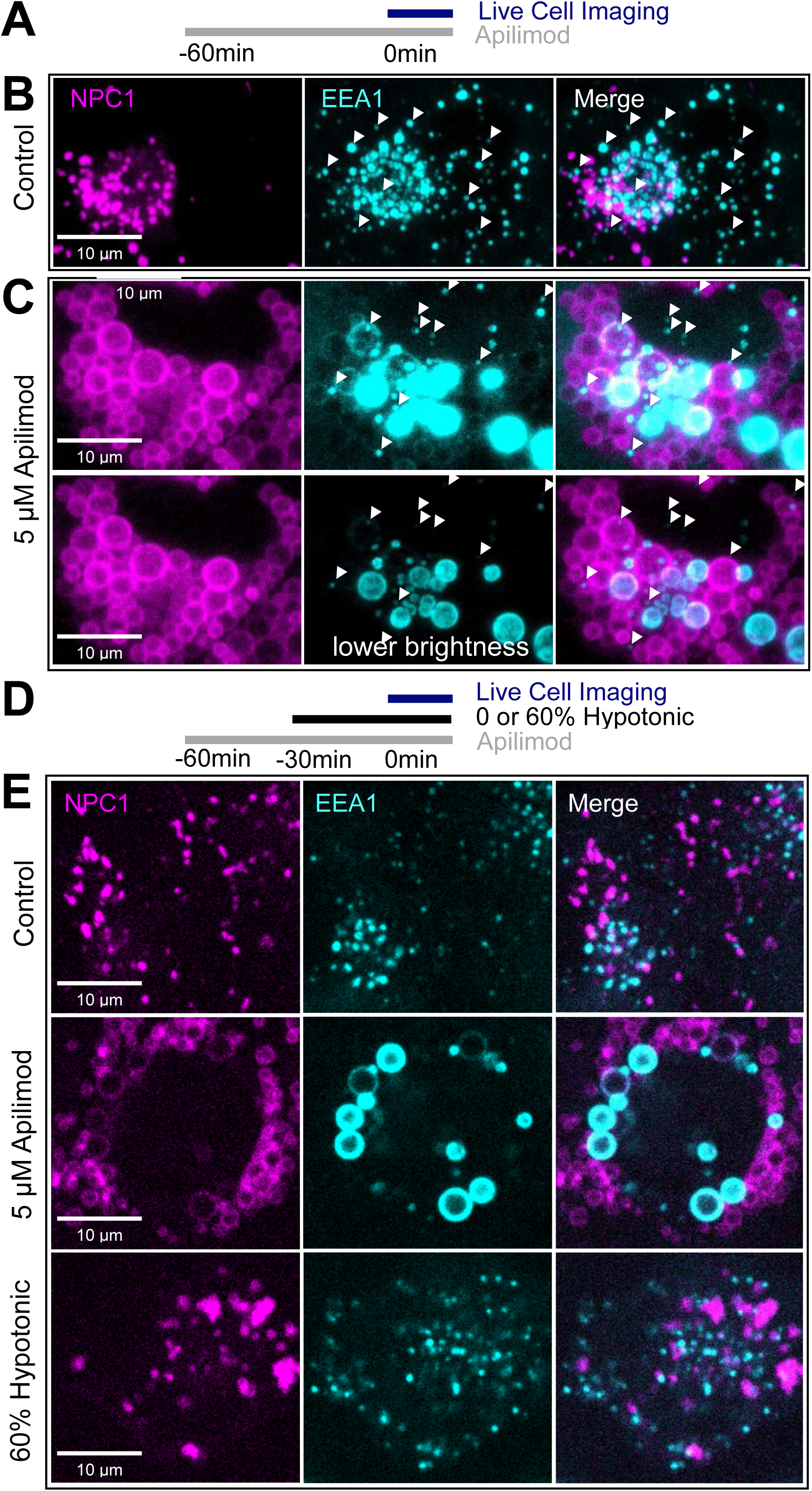
Apilimod fails to induce swelling of early endosomes EEA1-positive/NPC1-negative compartments (Related to Fig. 3). **(A)** Schematic of live cell imaging experiment. Gene-edited SVG-A cells expressing NPC1-Halo labeled with JFX 646 and EEA1-mScarlet were pretreated with 5 µM apilimod for 1 h and imaged live during a 10 min period, also in the presence of apilimod, using lattice light-sheet microscopy. **(B, C)** Representative maximum intensity z-projections in the absence (control **(B)**) or presence **(C)** of 5 µM apilimod. NPC1-Halo labeled with JFX 646 is shown in magenta, and EEA1-mScarlet in cyan. White arrowheads indicate early endosomes (EEA1-positive/NPC1-negative). Scale bar, 10 µm. **(C)** Top panel presents EEA1 images with higher contrast to visualize dimmer early endosomes; bottom panel presents the image in the same contrast as control **(B)**. **(D)** Schematic of the live-cell imaging experiment. Gene-edited SVG-A cells expressing NPC1–Halo (labeled with JFX646) and EEA1–mScarlet were pretreated with 5 µM apilimod for 30 min and imaged live by lattice light-sheet microscopy for 10 min in the continued presence of apilimod, under normal conditions or during incubation in 60% hypotonic medium (initiated 20 min before imaging and maintained throughout imaging). **(E)** Representative single plane images in absence or presence of 5 μM apilimod or 60% hypotonic media. NPC1-Halo labeled with JFX 646 is shown in magenta, and EEA1-mScarlet in cyan. Scale bar, 10 μm.

**Fig. S2.**
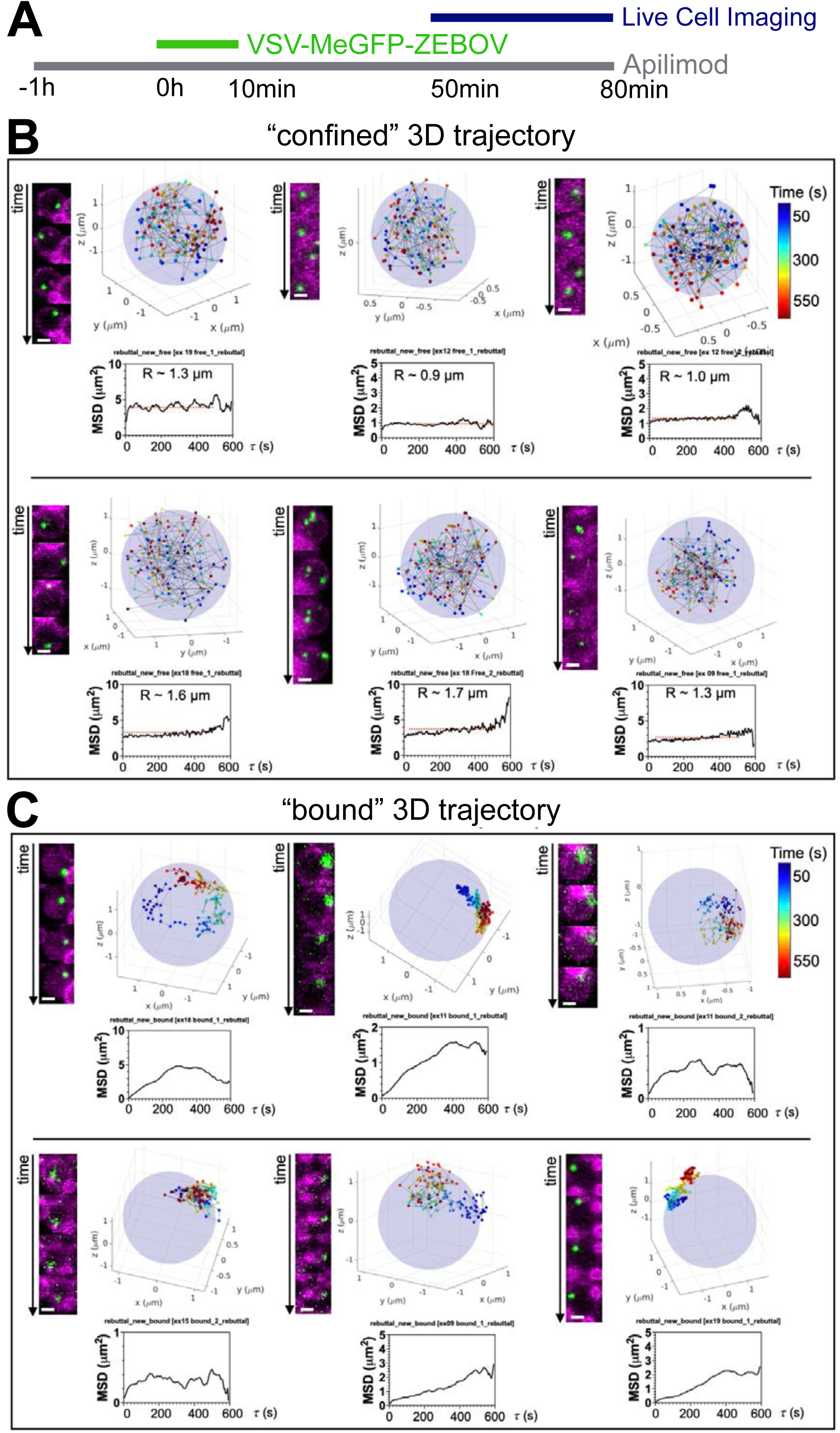
3D tracking of VSV-MeGFP–ZEBOV reveals confined virion populations within apilimod-induced swollen NPC1-positive endolysosomes. Associated with Fig 9 and Movies 1-3. **(A)** Schematic of the infection assay. Gene-edited SVG-A cells expressing NPC1-Halo were pretreated with 5 µM apilimod for 1 h, then incubated with VSV-MeGFP–ZEBOV (10 min, MOI ∼ 1) in the continued presence of 5 µM apilimod. Unbound virus was removed by three washes (each containing 5 µM apilimod), and cells were maintained in apilimod for the duration of the experiment. Live-cell imaging was performed at 50 min postinfection using lattice light-sheet microscopy. 3D time-lapse series were acquired every 4 s over 10 min. **(B, C)** Representative 3D trajectories of individual virions moving within the lumen **(B)** or along the limiting membrane **(C)** of NPC1-positive endolysosomes. NPC1-Halo–positive endolysosomes are shown as light blue circles. Virion positions are color-coded by time (dark blue to dark red). Insets show representative maximum-intensity z-projections from sequential images of an NPC1–Halo–positive endosome labeled with JF646 (magenta) and VSV-MeGFP–ZEBOV (green). Scale bar, 1 um. Mean square displacement (MSD) plots are shown; in **(B)**, the radius of confinement is derived from the MSD plateau using the relationship 6/5 R² (inset).

**Movies 1–3. Associated to Fig. 9 B and C.** Live-cell 3D time-lapse imaging of VSV-MeGFP–ZEBOV virions (green) associated with endolysosomes enlarged by PIKfyve inhibition with apilimod. SVG-A cells expressing gene-edited NPC1–Halo labeled with JF 646 (magenta). Cells were pretreated with 5 µM apilimod for 1 h and then incubated with fluorescent VSV-MeGFP–ZEBOV (10 min, MOI ∼1) in the continued presence of apilimod. Live-cell imaging by LLSM was performed 50 min postinfection. The 3D time-lapse series, acquired every 4 s for 10 min, are displayed as maximum-intensity projections. Scale bar, 2 µm.

